# Coarse-Grained Molecular Simulations and Ensemble-Based Mutational Profiling of Protein Stability in the Different Functional Forms of the SARS-CoV-2 Spike Trimers : Balancing Stability and Adaptability in BA.1, BA.2 and BA.2.75 Variants

**DOI:** 10.1101/2023.02.28.530489

**Authors:** Gennady Verkhivker, Mohammed Alshahrani, Grace Gupta

**Author notes:** Correspondence; (G.V).

## Abstract

The evolutionary and functional studies suggested that the emergence of the Omicron variants can be determined by multiple fitness trade-offs including the immune escape, binding affinity, conformational plasticity, protein stability and allosteric modulation. In this study, we embarked on a systematic comparative analysis of the conformational dynamics, electrostatics, protein stability and allostery in the different functional states of spike trimers for BA.1, BA.2, and BA.2.75 variants. Using efficient and accurate coarse-grained simulations and atomistic reconstruction of the ensembles, we examined conformational dynamics of the spike trimers that agrees with the recent functional studies, suggesting that BA.2.75 trimers are the most stable among these variants. A systematic mutational scanning of the inter-protomer interfaces in the spike trimers revealed a group of conserved structural stability hotspots that play a key role in modulation of functional dynamics and are also involved in the inter-protomer couplings through local contacts and interaction networks with the Omicron mutational sites. The results of mutational scanning provided evidence that BA.2.75 trimers are more stable than BA.2 and comparable in stability to BA.1 variant. Using dynamic network modeling of the S Omicron BA.1, BA.2 and BA.2.75 trimers we showed that the key network positions driving long-range signaling are associated with the major stability hotspots that are inter-connected along potential communication pathways, while sites of Omicron mutations may often correspond to weak spots of stability and allostery but are coupled to the major stability hotspots through interaction networks. The presented analysis of the BA.1, BA.2 and BA.2.75 trimers suggested that thermodynamic stability of BA.1 and BA.2.75 variants may be intimately linked with the residue interaction network organization that allows for a broad ensemble of allosteric communications in which signaling between structural stability hotspots may be modulated by the Omicron mutational sites. The findings provided plausible rationale for mechanisms in which Omicron mutations can evolve to balance thermodynamic stability and conformational adaptability in order to ensure proper tradeoff between stability, binding and immune escape.

## Introduction

The unprecedented body of structural and biochemical studies have explored mechanisms of SARS-CoV-2 infection showing the key role of the SARS-CoV-2 viral spike (S) glycoprotein undergoing stochastic movements between distinct functional forms [1–9]. The complex architecture of the S protein includes an amino (N)-terminal S1 subunit experiencing functional motions and structurally rigid carboxyl (C)-terminal S2 subunit. Conformational transformations of the SARS-CoV-2 S protein between the closed and open S states are exemplified by coordinated global movements of the S1 subunit consisting of an N-terminal domain (NTD), the receptor-binding domain (RBD), and two structurally conserved subdomains, SD1 and SD2, which together determine the structural and dynamic response of the S protein to binding partners and the host cell receptor ACE2 [10–15]. A number of biophysical studies provided in-depth characterization of the thermodynamic and kinetic aspects of the SARS-CoV-2 S functional trimer, showing the complex interplay between subdomain movements and long-range interactions that couple S1 and S2 subunits to modulate the RBD equilibrium and population-shifts between the RBD open (up) and closed (down) conformations regulating the exposure of the S protein to binding partners and strength of S-ACE2 binding [16–18]. The rapidly growing number of cryo-EM and X-ray structures of the SARS-CoV-2 S variants of concern (VOC’s) in various functional states and complexes with antibodies revealed a remarkable versatility of molecular mechanisms and diversity of binding epitopes that underlie binding affinity of S proteins with different classes of antibodies [19–28]. In particular, the cryo-EM structures of the S Omicron BA.1 variants in different functional states showed a subtle balance and tradeoffs of various factors driving binding thermodynamics in which mutations act cooperatively to regulate the open-closed equilibrium and promoting formation of the RBD-up states, to induce immune evasion by altering antibody epitopes, while simultaneously maintaining a structurally stable RBD-down state which allows for occlusion of highly immunogenic sites [27]. The enormous number of structural and functional studies of the SARS-CoV-2 S Omicron variants produced diverse and often conflicting hypotheses and rationales regarding the relative stability and functional role of the open and closed states in various Omicron variants. Several structural studies demonstrated that the S Omicron trimer can exhibit a much more compact architecture when compared to the S-Delta and native S-Wu-Hu-1 form [28]. The cryo-EM structures of the S Omicron BA.1 trimer unveiled in this study showed that the Omicron sites N856K, N969K, and T547K can promote favorable electrostatic interactions and lead to the hydrogen bonds with D658, Q755, and S982 from neighboring subunits, thus resulting in the increased number of the inter-protomer contacts which can confer the enhanced stability for the Omicron S-trimer [28]. At the same time, differential scanning calorimetry experiments indicated that the folding stability of the S Omicron BA.1 and S Wu-Hu-1 trimers is very similar [29]. The biophysical analysis of protein stability for the Wu-Hu-1, Delta and Omicron variants was also done by using differential scanning fluorimetry (DSF) assay that measured the inflection temperature and revealed that the transition to the folding state for the S Omicron BA.1 was shifted to lower temperatures as compared to the S Wu-Hu-1 and S Delta, suggesting the reduced protein stability of the S Omicron BA.1 variant [29,30]. The S Omicron BA.1 and BA.2 S trimers were shown to adopt stable RBD-down trimer conformations that are stabilized by a strong network of the inter-protomer contacts, leading to the higher thermostability [31,32]. The thermostability of the S-D614G, S-BA.1, and SB-BA.2 protein ectodomains as well as the stability of their corresponding monomeric RBD constructs were evaluated in the DSF assays showing the reduced stability of the BA.1 RBD, while BA.2 RBD appeared to be more stable than BA.1 but less stable than the Wu-Hu-1 [31]. The reduced stability of the BA.1 RBD relative to the native Wu-Hu-1 RBD is also in agreement with other published reports [32]. The important and somewhat unexpected revelation of some structural studies was that S Omicron BA.1 trimer may preferentially adopt the 1RBD-up conformation both before and after ACE2 binding where the S371L, S373P and S375F substitutions tend to enhance the stability of the 1RBD-up conformation and prevent exposure of more up-RBDs to ACE2 binding [33]. Similar results were obtained in other structural investigation of the Omicron BA.1 variant, presenting evidence of the stabilization of the open S conformation due to the enhanced inter-domain and inter-subunit packing, induced by mutations of the Omicron strain [28,34,35]. The reported 3.1 Å-resolution cryo-EM structure of the S Omicron protein ectodomain [34] and another 3.0 Å cryo-EM structure of the S Omicron protein ectodomain [35] showed that in contrast to the original strain of SARS-CoV-2 with a mixture of open and closed conformations, the S Omicron BA.1 proteins may adopt predominantly an open 1 RBD-upright position predisposed for receptor binding.

Structural studies of the Omicron variant indicated that evolutionary pressure invokes a complex interplay of thermodynamic factors between mutations that increase affinity for the ACE2 with other RBD modifications that disfavor ACE2 binding but facilitate immune escape [36–40]. These investigations underscored a mechanism that involves balancing ACE2 affinity enhancement with the need to produce immune-escaping mutations. According to these studies T478K, Q493R, G496S, and Q498R strengthened the binding of Omicron to ACE2 while mutations K417N and E484A decreased the binding affinity. The cryo-EM study of the S Omicron BA.1 variant examined binding and antigenic properties by bio-layer interferometry showing that the improved S-ACE2 binding may be attributed largely to the effect of N501Y, Q493R and Q498R mutations [41]. At the same time, Omicron mutations S477N, T478K, E484A and K417N may induce a moderate loss in ACE2 binding while promoting the increased neutralization escape potential of the Omicron variant from antibodies [41]. Atomic force microscopy studies revealed the role of mechanical stability in mediating immune evasion, pointing to a combined effect of mechanical forces, protein stability and binding interactions that collectively control the virus fitness advantage and immune escape mechanisms [42–44]. The structural basis of the higher binding affinity of ACE2 to currently circulating Omicron subvariants was elucidated in an investigation which reported the structures of the RBD-ACE2 complexes for BA.1.1, BA.2, and BA.3 variants [45]. This study showed that the Omicron BA.1.1 and BA.2 binding affinities with ACE2 are stronger than binding for BA.3 and BA.1 subvariants. The enhanced transmission of Omicron BA.2 was examined using structural and biochemical analysis of binding with the human ACE2 (hACE2) showing that the S Omicron BA.2 trimer displayed ACE2 binding affinity which is 11-fold higher than that of the S Wu-Hu-1 trimer and 2-fold higher than that of the S Omicron BA.1 spike [46].

The recent biophysical study employed surface plasmon resonance (SPR) tools to measure the binding affinity of the Omicron BA.4/5 RBD for ACE2 which was stronger as compared with the ancestral Wu-Hu-1 strain, BA.1, and BA.2 (3-, 3-, and 2-fold, respectively) [47]. An impressive structural and biochemical study reported cryo-EM structures of the S trimers for BA.1, BA.2, BA.3, and BA.4/BA.5 subvariants of Omicron [48]. The binding affinities of the Omicron variants with the ACE2 receptor determined by SPR showed a decreased binding affinity for the BA.4/BA.5 subvariants, also revealing that the Omicron BA.2 displayed slightly higher binding affinities than the other Omicron variants. The cryo-EM structures showed that BA.1 S-trimer is stabilized in an open conformation, BA.2 exhibits two conformational states corresponding to a closed and an open form with one RBD in the up position, while S BA.3 and S BA.4 trimers adopt closed or semi-closed forms [48]. The thermal stability assays employed in this illuminating study also verified that S-trimers from BA.2 sublineages (B.2, B.2.12.1) were the least stable among B.1, B.2, B.3 and B.4 variants. According to this study, S-trimers from the BA.2 sublineage displayed a somewhat less compact architecture while BA.1, BA.3 and BA.4/BA.5 spike exhibited tighter inter-subunit organization with more buried areas between S2 subunits [48]. Structure-functional studies of the Omicron BA.1, BA.2, BA.2.12.1, BA.4 and BA.5 subvariants showed the increased ACE2 binding affinity and stronger evasion of neutralizing antibody responses for these Omicron variants as compared to the Wu-Hu-1 and Delta strains, confirming that the compounded effect of the enhanced ACE2 receptor binding and stronger immune evasion may have contributed to the rapid spread of these Omicron sublineages [49]. Using bio-layer interferometry and SPR techniques, this investigation also quantified and confirmed the greater binding affinity of the S Omicron BA.2 as compared to the native Wu-Hu-1 strain and original Omicron BA.1 variant [49]. Recent structural studies reported cryo-EM conformations of the BA.2.75 S trimer in the open and closed forms as well as structures of the open BA.2.75 S trimer complexes with ACE2 [50]. This study also revealed in viro thermal stabilities of the Omicron variants at neutral pH, showing that the BA.2.75 S-trimer was the most stable, followed by BA.1, BA.2.12.1, BA.5 and BA.2 variants. The binding affinities between hACE2 and RBDs were evaluated the six Omicron subvariants (BA.1, BA.2, BA.3, BA.4/5, BA.2.12.1, and BA.2.75), together with the other earlier four VOCs (Alpha, Beta, Gamma, and Delta) by SPR revealing that BA.2.75 displayed 4-6-fold increased binding affinity to hACE2 compared with other Omicron variants [50]. Remarkably, this pioneering study indicated that the structural heterogeneity of BA.2.75 S trimer can coexist with the increasingly stable open conformation featuring a more rigid and compact RBD conformations than other subvariants, which may be one of the contributing factors underlying the increased thermal stability of the S BA.2.75 trimer [50]. Importantly, this study confirmed that BA.2.75 has the highest ACE2 affinity among all SARS-CoV-2 variants with the known experimental binding measurements. The role of N460K mutation in binding is unclear as this residue resides outside of the binding epitope. It is possible that R493Q and N460K may cooperate with other mutational sites to increase binding. An antigenic and biophysical characterization of BA.2.75 variant unveiled the better balance between immune evasion and ACE2 binding, where the binding affinity to ACE2 is increased 9-fold as compared to the BA.2 variant arguably owing to N460K and reversed R493Q mutations [51]. It was conjectured that these mutations may marginally affect neutralization escape and prioritize ACE2 binding affinity to increase the transmissibility of BA.2.75 [51]. However, this rationale has not been rigorously validated and remains a working hypothesis that requires further confirmation. According to the latest structure-functional investigation of BA.2.75 variant [52] NTD mutations K147E, F157L, and I210V, and two substitutions in the RBD, N460K and R493Q, significantly increased infectivity, which is particularly striking for N460K substitution increasing infectivity by 44-fold. At the same time, it was established that BA.2.75 variant can be endowed with significant antibody evasion properties and greater ACE2 binding as well as improved growth efficiency and intrinsic pathogenicity over BA.2 variant. The receptor-binding assay performed in this study showed that G446S significantly decreased the binding affinity of BA.2 RBD to ACE2 and was acquired to evade antiviral immunity, while other substitutions in the BA.2.75 S RBD, particularly N460K may compensate for this loss of binding affinity [52]. Another insightful study showed that G446S and N460K mutations of BA.2.75 are primarily responsible for its enhanced resistance to neutralizing antibodies, while R493Q reversion reduces BA.2.75 neutralization resistance [53]. According to this study, BA.2.75 shows enhanced cell-cell fusion over BA.2, driven largely by the N460K mutation, which enhances S processing. Similar mechanistic conclusions were reached in a study focusing on Omicron BA.5 [54] showing that the R493Q reversion in the BA.4/5 S protein potentially contributes to evading immunity and the effect of the R493Q on attenuating the resistance could be canceled by F486V substitution while L452R compensated the decreased ACE2 binding affinity. These studies suggested that the acquisition of functionally balanced substitutions where some mutations promote immune evasion, but tend to decrease ACE2 affinity, while the others can result in the increased ACE2 affinity to compensate for the effect of immune-resistant modifications might be a common strategy of SARS-CoV-2 evolution shared by Omicron subvariants. The newly emerging variants such as BA.2.3.20, BA.2.75.2, BQ.1.1 and especially XBB, a recombinant of BJ.1 and BM.1.1.1 display substantial growth advantages over previous Omicron variants, and some RBD residues including R346, K356, K444, V445, G446, N450, L452, N460, F486, F490, R493 and S494 were found mutated in at least five independent Omicron sublineages that exhibited a high growth advantage [55]. By examining forces driving the accelerated emergence of convergent RBD mutations it was suggested that the immune pressure on the RBD becomes increasingly focused and promotes convergent evolution, explaining the observed sudden acceleration of SARS-CoV-2 RBD evolution and the convergence pattern [55].

Computer simulations provided important atomistic and mechanistic advances into understanding the dynamics and function of the SARS-CoV-2 S proteins. All-atom molecular dynamics (MD) simulations of the full-length SARS-CoV-2 S glycoprotein embedded in the viral membrane, with a complete glycosylation profile provided detailed characterization of the conformational landscapes of the S proteins in the physiological environment [56–61]. Using distributed cloud-based computing, large scale MD simulations of the viral proteome observed dramatic opening of the S protein complex, predicting the existence of several cryptic epitopes in the S protein [62]. MD simulations of the S-protein in solution and targeted simulations of conformational changes between the open and closed forms revealed the key electrostatic interdomain interactions mediating the protein stability and kinetics of the functional spike states [63]. Using the replica-exchange MD simulations with solute tempering of selected surface charged residues, the conformational landscapes of the full-length S protein trimers were investigated, unveiling the transition pathways via inter-domain interactions, hidden functional intermediates along open-closed transition pathways and previously unknown cryptic pockets [64]. The enhanced MD simulations examined the stability of the S-D 614G variant showing that the mutation orders the 630-loop structure and allosterically alters global interactions between RBDs, forming an asymmetric and mobile down conformation and facilitating transitions toward up conformation [65]. Our previous studies revealed that the SARS-CoV-2 S protein can function as an allosteric regulatory machinery that can exploit the intrinsic plasticity of functional regions controlled by stable allosteric hotspots to modulate specific regulatory and binding functions [66–72]. A number of computational studies employed atomistic simulations and binding energy analysis to examine the interactions between the S-RBD Omicron and the ACE2 receptor. All-atom MD simulations of the S Omicron trimer and the Omicron RBD–ACE2 complexes suggested that the Omicron mutations may have evolved to inflict a greater infectivity using a combination of more efficient RBD opening, the increased binding affinity with ACE2, and optimized capacity for antibody escape [73]. MD simulations of the Omicron RBD binding with ACE2 suggested that K417N, G446S, and Y505H mutations can decrease the ACE2 binding, while S447N, Q493R, G496S, Q498R, and N501Y mutations improve binding affinity with the host receptor [74]. By examining a large number of mutant complexes, it was found that high-affinity RBD mutations tend to cluster near known human ACE2 recognition sites supporting the view that combinatorial mutations in SARS-CoV-2 can develop in sites amenable for non-additive enhancements in binding and antibody evasion. simultaneously maintain high-affinity binding to ACE2 and evade antibodies (e.g., by N440K, L452R, E484K/Q/R, K417N/T) [75]. We examined differences in allosteric interactions and communications in the S-RBD complexes for Delta and Omicron variants using a combination of perturbation-based scanning of allosteric propensities and dynamics-based network analyses [76]. This study showed that G496S, Q498R, N501Y and Y505H correspond to the key binding energy hotspots and also contribute decisively to allosteric communications between S-RBD and ACE2. It was also suggested that SARS-CoV-2 S protein may exploit plasticity of the RBD to generate escape mutants while engaging a small group of functional hotspots to mediate efficient local binding interactions and long-range allosteric communications with ACE2 [76]. All-atom MD simulations of the RBD-ACE2 complexes for BA.1 BA.1.1, BA.2, an BA.3 Omicron subvariants were combined with a systematic mutational scanning of the RBD-ACE2 binding interfaces, revealing functional roles of the key Omicron mutational site R493, R498 and Y501 acting as binding energy hotspots, drivers of electrostatic interactions and mediators of epistatic effects and long-range communications [77].

The evolutionary and functional studies suggested that the emergence of the Omicron variants can be determined by multiple fitness trade-offs including the immune escape, binding affinity for ACE2, conformational plasticity, protein stability and allosteric modulation [78–80]. To investigate and quantify this fundamental hypothesis, we embarked on a systematic comparative analysis of the conformational dynamics, protein stability and allostery in the different functional states of S trimers for BA.1, BA.2, and BA.2.75 variants. In the current study, we combined multiple coarse-grained molecular simulations followed by atomistic reconstruction of the trajectories, electrostatic analysis, a comprehensive mutational scanning using two different approaches and ensemble-based dynamic network analysis to simulate both 1RBD-up open and closed states of the S trimers for BA.1, BA.2 and BA.2.75 Omicron variants. Using this battery of computational approaches, we dissect important dynamic and energetic patterns underlying the inter-protomer interactions as well as identify the protein stability hotspots in the S Omicron BA.1, BA.2 and BA.2.75 trimer. The results show that BA.2.75 variant can be characterized by the greater stability among the studied Omicron variants, while BA.2 trimers are characterized by a considerable mobility and are the least stable. Our findings agree with the latest experimental data and provide an important rationale behind the enhanced stability of the BA.2.75 trimers as one of the factors driving its increased infectivity and binding. Through a detailed analysis of the dynamics and intermolecular interactions, we characterize the fundamental commonalities and differences in the organization and dynamics of the S Omicron BA.1, BA.2 and BA.2.75 trimers. Our results suggest that Omicron sites are coupled to major stability hotspots through local and long-range interactions allowing for modulation of stability and dynamic changes in the S protein without compromising the integrity and intrinsic activity of the spike. These results further highlight the underlying mechanism of S proteins that is linked with the subtle balancing and tradeoff between protein stability, conformational adaptability and plasticity that collectively drive evolution of the SARS-CoV-2 S variants. Using conformational ensembles and dynamic network modeling of the S Omicron BA.1, BA.2 and BA.2.75 trimers we show that the key network positions driving long-range signaling are associated with the major stability hotspots that are inter-connected along potential communication pathways, while sites of Omicron mutations in S1 subunit may correspond to weak spots of stability and allostery but are coupled to the major regulatory positions through interaction networks. We argue that this allows SARS-CoV-2 virus to evolve mutations in these positions without compromising protein stability and spike viability while allowing for modulation of allosteric interactions and conformational changes.

## 2. Results and Discussion

### Coarse-Grained Molecular Simulations s Reveal Common and Distinct Signatures of Conformational Stability and Flexibility in the SARS-CoV-2 S Omicron Variants

To examine structural and dynamic signatures of the Omicron variants, we embarked on a comparative analysis of the conformational landscapes for S Omicron BA.1. BA.2 and BA.2.75 trimers in the closed forms (Figure 1) and open forms (Figure 2) using coarse-grained Brownian dynamics (CG-BD) simulations within the ProPHet (Probing Protein Heterogeneity) approach [81–84]. Due to large size of the S trimers, we opted to carry out independent CG-BD simulations of the S Omicron BA.1, BA.2 and BA.2.75 trimer structures followed by subsequent atomistic reconstruction which allowed for subsequent detection of reproducible dynamic patterns and subtle differences in the intrinsic dynamics induced by Omicron subvariants. A total of 12 mutations (G339D, S373P, S375F, K417N, N440K, S477N, T478K, E484A, Q493R, Q498R, N501Y, and Y505H) are shared among the BA.1 and BA.2 variants. In the RBD, BA.1 contains unique mutations S371L, G446S, and G496S while BA.2 carries S371F, T376A, D405N, and R408S mutations (Table 1). Structural analysis of the RBD binding epitopes in the homotrimers and complexes with ACE2 (Table 2) revealed a very similar composition of the interacting residues for all studied Omicron subvariants and virtually identical topography of the binding interface (Figures 1,2). BA.2.75 variant has nine additional mutations as compared to its parent BA.2 including NTD (K147E, W152R, F157L, I210V, and G257S) and RBD (D339H, G446S, N460K, and R493Q) (Table 1) [85]. Several BA.2.75 mutations are shared with other Omicron sublineages. G446S mutation is shared with Omicron BA.1, and R493Q reversion is present in BA.4/BA.5. It was established that BA.2.75 variant can be endowed with significant antibody evasion properties and greater ACE2 binding over BA.2 variant, largely attributed to acquisition of N460K and reverse R493Q mutations in the RBD [52,85].

**Figure 1.**
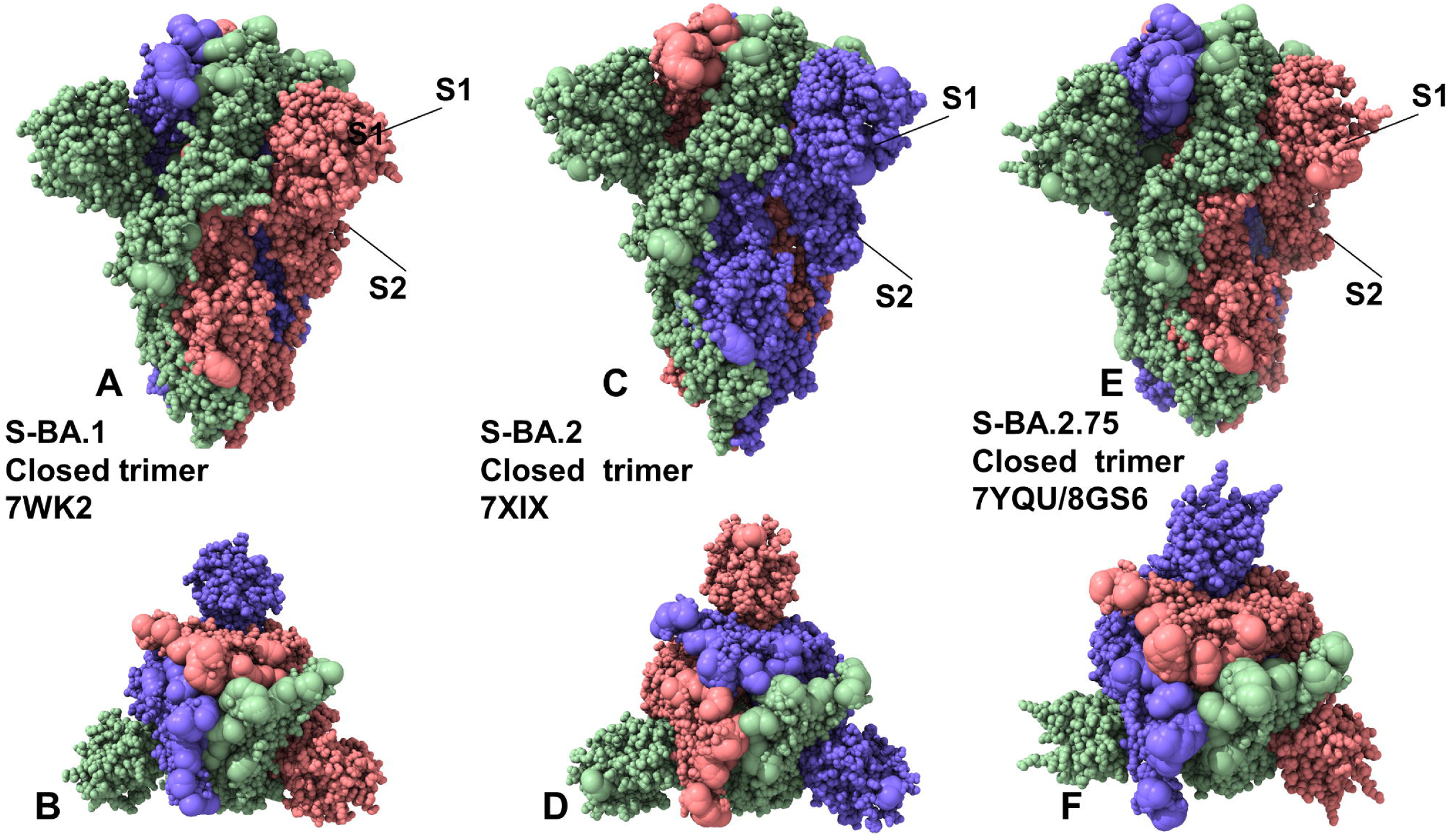
Structural organization and mapping of the Omicron mutations in the SARS-CoV-2 S closed Omicron BA.1, BA.2 and BA2.75 trimers. (A) The cryo-EM structure of the S Omicron BA.1 trimer in the closed 3RBD-down form (pdb id 7WK2). (B) The top view of the closed BA.1 trimer. The S trimer is shown in full spheres and colored by the protomer. The Omicron BA.1 sites (A67, T95I, G339D, S371L, S373P, S375F, K417N, N440K, G446S, S477N, T478K, E484A, Q493R, G496S, Q498R, N501Y, Y505H, T547K, D614G, H655Y, N679K, P681H, N764K, D796Y, N856K, Q954H, N969K, L981F) are shown in larger spheres. (C) The cryo-EM structure of the S Omicron BA.2 trimer in the closed, 3 RBD-down form (pdb id 7XIX). (D) The top view of the closed BA.2 trimer. The Omicron BA.2 sites (T19I, G142D, V213G, G339D, S371F, S373P, S375F, T376A, D405N, R408S, K417N, N440K, S477N, T478K, E484A, Q493R, Q498R, N501Y, Y505H, D614G, H655Y, N679K, P681H, N764K, D796Y, Q954H, N969K) are shown in large spheres. (E) The cryo-EM structure of the S Omicron BA.2.75 trimer in the closed, 3 RBD-down form (pdb id 7YQU/8GS6). (D) The top view of the closed BA.2.75 trimer. (F) The top view of the closed S Omicron BA.2.75 trimer. The Omicron BA.2.75 sites (T19I, G142D, K147E, W152R, F157L, I210V, V213G, G257S, G339H, S371F, S373P, S375F, T376A, D405N, R408S, K417N, N440K, G446N, N460K, S477N, T478K, E484A, Q498R, N501Y, Y505H, D614G, H655Y, N679K, P681H, N764K, D796Y, Q954H, N969K) are shown in larger spheres.

**Figure 2.**
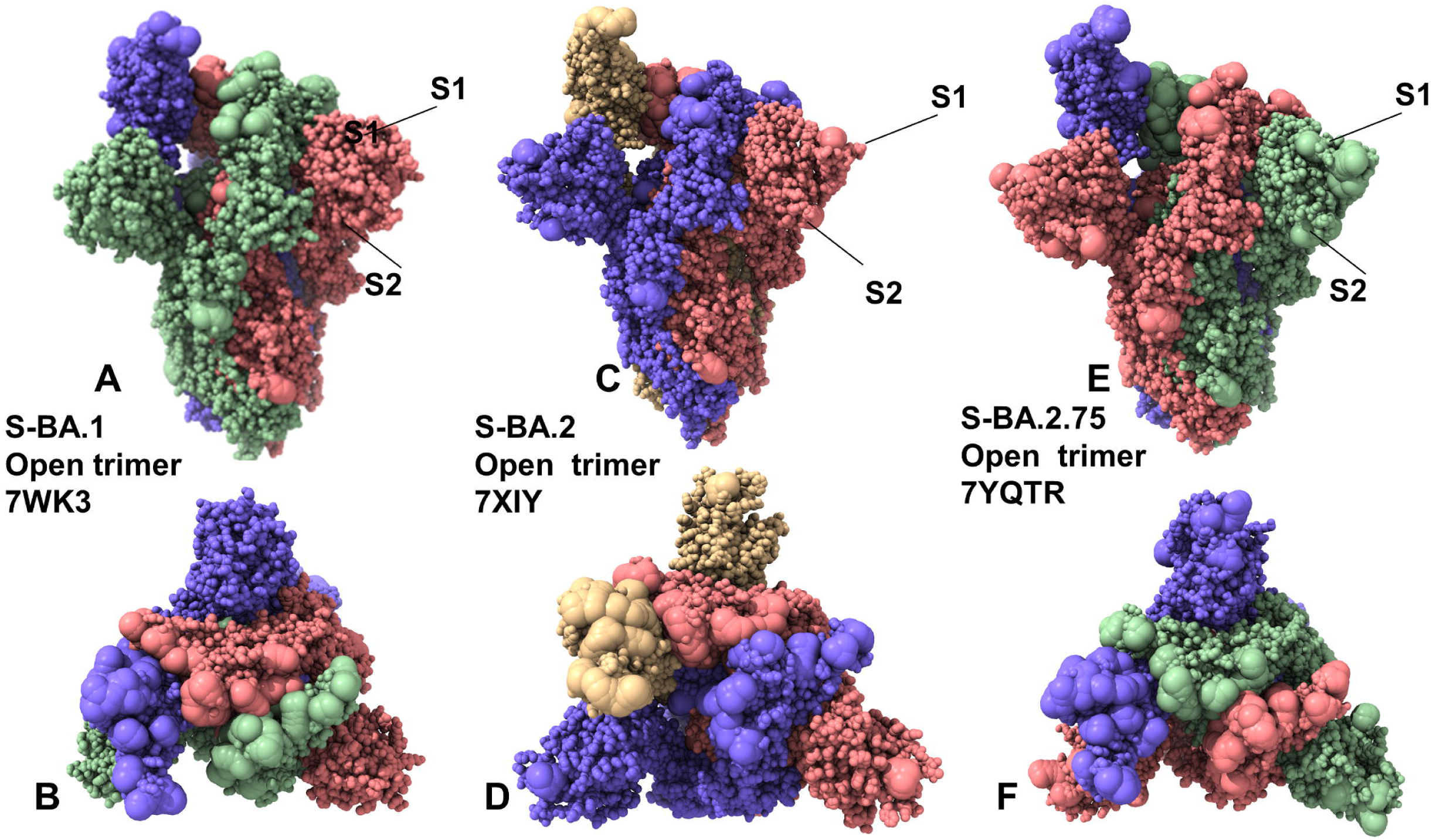
Structural organization and mapping of the Omicron mutations in the SARS-CoV-2 S Omicron BA.1, BA.2 and BA2.75 open trimers. (A) The cryo-EM structure of the S Omicron BA.1 trimer in the open 1RBD-up form (pdb id 7WK3). (B) The top view of the open BA.1 trimer. The S trimer is shown in full spheres and colored by the protomer. The Omicron BA.1 mutational sites are shown in larger spheres. (C) The cryo-EM structure of the S Omicron BA.2 trimer in the open 1RBD-up (pdb id 7XIW). (D) The top view of the closed BA.2 trimer. The Omicron BA.2 mutational sites are shown in large spheres. (E) The cryo-EM structure of the S Omicron BA.2.75 trimer in the open 1RBD-up form (pdb id 7YQT). (D) The top view of the closed BA.2.75 trimer. (F) The top view of the closed S Omicron BA.2.75 trimer. The Omicron BA.2.75 sites are shown in larger spheres.

**Table 1.** Mutational landscape of the Omicron mutations for BA.1, BA.2 and BA.275 variants

| **SARS-CoV-2 Omicron Variant** | **Mutational landscape of the RBD** |
| --- | --- |
| BA.1 | A67, T95I, G339D, S371L, S373P, S375F, K417N, N440K, G446S, S477N, T478K, E484A, Q493R, G496S, Q498R, N501Y, Y505H, T547K, D614G, H655Y, N679K, P681H, N764K, D796Y, N856K, Q954H, N969K, L981F |
| BA.2 | T19I, G142D, V213G, G339D, S371F, S373P, S375F, T376A, D405N, R408S, K417N, N440K, S477N, T478K, E484A, Q493R, Q498R, N501Y, Y505H, D614G, H655Y, N679K, P681H, N764K, D796Y, Q954H, N969K |
| BA.2.75 | T19I, G142D, K147E, W152R, F157L, I210V, V213G, G257S, G339H, S371F, S373P, S375F, T376A, D405N, R408S, K417N, N440K, G446N, N460K, S477N, T478K, E484A, Q498R, N501Y, Y505H, D614G, H655Y, N679K, P681H, N764K, D796Y, Q954H, N969K |

**Table 2.** Structures of the S Omicron BA.2 and BA.2.75 variants.

| Variant | RBDs | Ligand | Resolution | PDB |
| --- | --- | --- | --- | --- |
| BA.2 | 3 down | - | 3.25 | 7xix |
| BA.2 | 3 down | - | 3.31 | 7ub0 |
| BA.2 | 3 down | - | 3.35 | 7ub5 |
| BA.2 | 3 down | - | 3.52 | 7ub6 |
| BA.2 | 1 up | - | 3.62 | 7xiw |
| BA.2 | 1 up | 1 ACE-2 | 3.20 | 7xoa |
| BA.2 | 2 up | 2 ACE-2 | 3.30 | 7xob |
| BA.2 | 2 up | 2 ACE-2 | 3.38 | 7xo7 |
| BA.2 | 3 up | 3 ACE-2 | 3.48 | 7xo8 |
| BA.2.75 | 3 down | - | 2.86 | 8gs6 |
| BA.2.75 | 3 down | - | 3.19 | 7yqu |
| BA.2.75 | 3 down | - | 3.51 | 7yqw |
| BA.2.75 | 1 up | - | 3.45 | 7yqt |
| BA.2.75 | 1 up | - | 3.58 | 7yqv |
| BA.2.75 | 1 up | 1 ACE-2 | 3.30 | 7yr2 |
| BA.2.75 | 2 up | 2 ACE-2 | 3.52 | 7yr3 |

Multiple cryo-EM structures of the S Omicron trimers were used in CG-BD simulations (Table 3) which allowed for a comparative analysis of protein dynamics in the distinct S Omicron states. Although the magnitude of the protein residue fluctuations derived from CG-BD simulations may be affected approximate coarse-grained nature of the energetic force field, the ensemble-averages of these simplified trajectories yielded a robust differentiation of rigid and flexible regions, also pointing to the intrinsic flexibility patterns of the open and closed states. We analyzed the root mean square fluctuation (RMSF) profiles obtained from CG-BD simulations of the S Omicron trimer in the closed state with all 3 RBDs in the down conformation (Figure 3). Despite generally similar shapes of the RMSF profiles for all Omicron subvariants, which may be expected given a high degree of structural similarity of the closed trimer, we also observed important dynamic differences.

**Table 3.** Structures of the S Omicron trimers examined in this study.

| **PDB** | **System** | **Per simulation** | **# simulations** |
| --- | --- | --- | --- |
| 7WK2 | Omicron BA.1 closed trimer | 500,000 steps | 100 |
| 7WK3 | Omicron BA.1 open timer | 500,000 steps | 100 |
| 7XIX | Omicron BA.2 closed trimer | 500,000 steps | 100 |
| 7XIW | Omicron BA.2 open trimer | 500,000 steps | 100 |
| 7YQU | Omicron BA.2.75 closed trimer | 500,000 steps | 100 |
| 8GS6 | Omicron BA.2.75 closed trimer | 500,000 steps | 100 |
| 7YQT | Omicron BA.2.75 open trimer | 500,000 steps | 100 |

**Figure 3.**
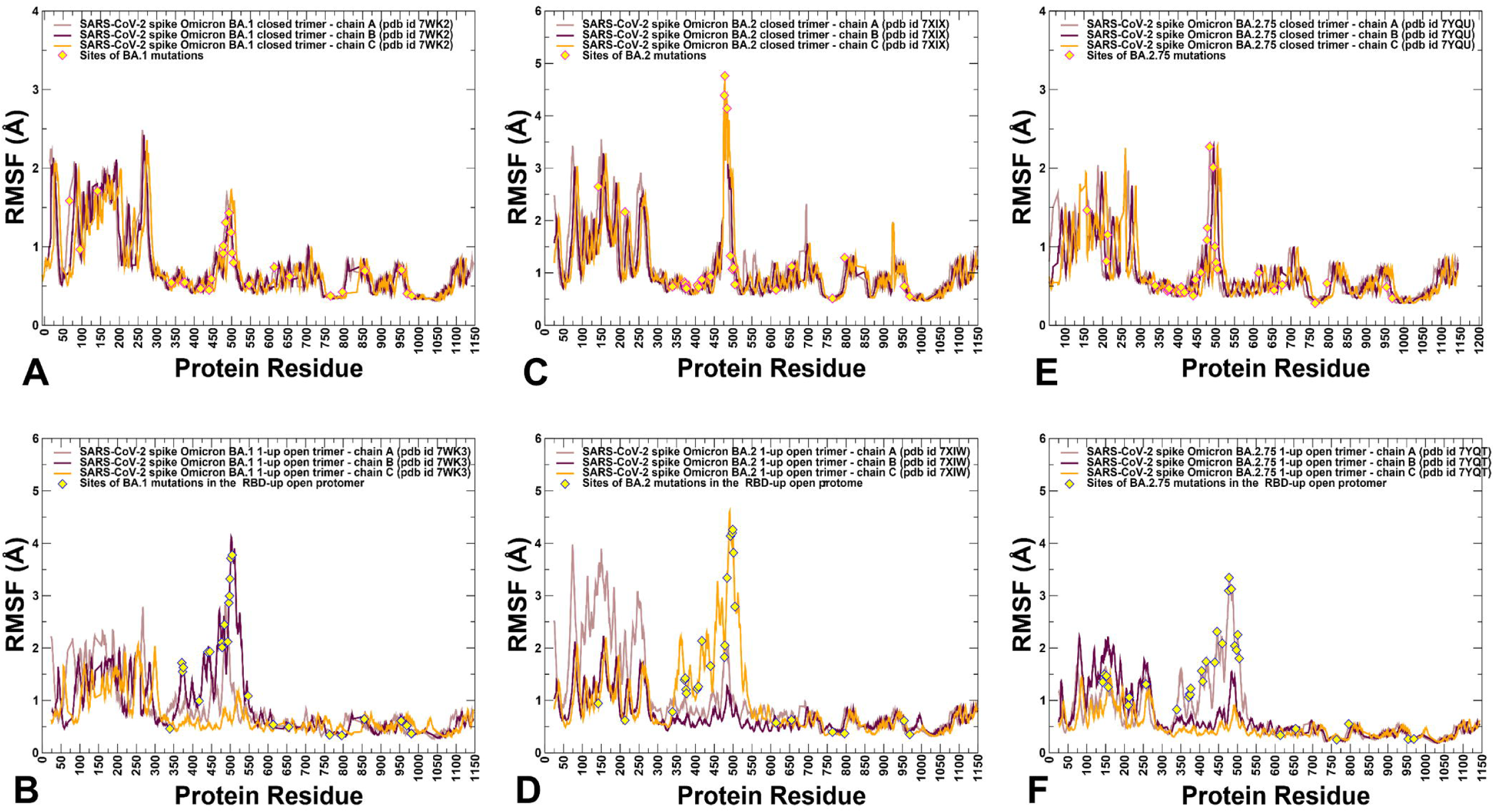
Conformational dynamics profiles of the SARS-CoV-2 S trimers. (A) The root mean square fluctuations (RMSF) profiles for the S Omicron BA.1 closed trimer (pdb id 7WK2). The RMSFs for protomer A are in brown lines, for protomer B in maroon lines, and for protomer C in orange lines. (B) The RMSF profiles for the S Omicron BA.1 trimer in the open form (pdb id 7WK3). The 1RBD-up protomer in this structure is protomer A shown in brown lines. The Omicron BA.1 mutations are shown yellow-filled diamonds. (C) The RMSF profiles for the SARS-CoV-2 S Omicron BA.2 closed trimer (pdb id 7XIX). The RMSFs for protomer A are in brown lines, for protomer B in maroon-colored lines, and for protomer C in orange lines. (D) The RMSF profiles for the S Omicron BA.2 trimer in the open form (pdb id 7XIW). The 1RBD-up protomer in this structure is protomer C shown in orange lines. The Omicron BA.2 mutations are in yellow-filled diamonds. (E) The RMSF profiles for the SARS-CoV-2 S Omicron BA.2.75 closed trimer (pdb id 7YQU). The RMSFs for protomer A are in brown lines, for protomer B in maroon-colored lines, and for protomer C in orange lines. (F) The RMSF profiles for the S Omicron BA.2.75 trimer in the open form (pdb id 7YQT). The 1RBD-up protomer in this structure is protomer A shown in brown lines. The positions of the Omicron BA.2.75 mutations are shown yellow-filled diamonds.

In the Omicron BA.1 closed trimer, relatively high fluctuations in the NTD regions can be contrasted to small moderate movements in the S1 domain, including the RBD regions as well as small displacements for the S2 subdomain (Figure 3A). The upstream helix (UH) (residues 736-781) and central helix (CH) (residues 986-1035) are particularly rigid in the S2 domain, while CTD1 (residues 528-591) and CTD2 (residues 592-686) undergo only moderate fluctuations (Figure 3A). The positions of the BA.1 mutations in the RBD of the closed trimer are associated with some fluctuations, particularly for S477N, T478K, Q493R, Q498R mutations. However, other important BA.1 RBD mutations N501Y and Y505H showed considerable rigidity and are stabilized by the closed trimer arrangement of the RBD-RBD packing (Figure 3A). In contrast, for BA.2 closed trimer, the thermal fluctuations of the NTD and RBD regions are appreciably larger, while the S2 regions remain stable (Figure 3C). In the BA.1 N856K enables formation of salt-bridge and hydrogen bond with the residues D568 and T572, respectively on the neighboring protomer while N764K forms a hydrogen bond with residue N317 on the neighboring protomer and salt-bridge with residue D737 on the same protomer. The increased rigidity of these residues in simulations suggested these interactions could contribute to stabilization of the BA.1 closed trimer. In the S BA.2 trimer, the overall degree of mobility is markedly increased as compared to the S Omicron BA.1. These findings are consistent with the experimental evidence revealing the increased heterogeneity of the S protein induced by BA.2 mutations and the decreased stability [48,50]. Consistent with the structural and biophysical data, CG-BD simulations revealed the increased stabilization of the closed S BA.2.75 trimer (Figure 3E). We observed that BA.2.75 mutational positions in the RBD remained stable and only S477N and T478K residues experienced some appreciable deviations (Figure 3E). The RBD mutational sites G339H, S371F, S373P, S375F showed considerable rigidity while N460K mutation did not increase mobility of the RBD loop and may even improve stabilization in this region (Figure 3E). It appeared that N460K residue forms a stable intramolecular salt bridge with D420, leading to the reduced mobility of the distal loop and supporting the notion that N460K may contribute to the RBD folding and the overall stability of the S BA.2.75 closed trimer (Figure 3C). According to the experimental data S-BA.2 spike exhibits two conformational states corresponding to a closed form, with all three RBDs in the down configuration and an open form with one RBD in the up position [48]. Conformational dynamics showed that the closed states may be more stable in the S BA.1 and S-BA.2.75 closed trimers as compared to the S BA.2 (Figure 3).

CG-BD simulations of the 1RBD-up open trimers for Omicron subvariants provided a more significant differentiation of the protein dynamics. Importantly, we found that the BA.2 open trimer (Figure 3D) is considerably more flexible than BA.1 (Figure 3B) and BA.2.75 open conformations (Figure 3F). As may be expected, the RBD-up conformation experienced significant fluctuations, but additionally we observed larger displacements of the NTD regions in the down protomer of the open BA.2 trimer (Figure 3D). Considerably smaller fluctuations were seen in simulations of BA.1 open trimer (Figure 3B), showing appreciable displacements for the RBD-up and also for one of the RBD-down conformations. In the RBD-up protomer, Omicron mutational sites N440K, G446S, S477N, T478K, E484A, Q493R, G496S, Q498R and N501Y experienced noticeable degree of mobility, while the NTD regions remained relatively stable. The most interesting differences were seen in the dynamics profile of the BA.2.75 open trimer, showing very small deviations for the two RBD-down protomers and suppressed mobility of the RBD-up protomer (Figure 3F) as compared to the other open trimers. Interestingly, mapping of the BA.2.75 mutational sites onto the dynamics profile showed that most of these positions including G339H and N460K displayed only modest RMSF values < 2.0 Å and remained mostly stable in the course of simulations. Only S477N, T478K and E484A positions showed RMSFs ∼ 2.8-3.0 Å while other important sites R493Q, Q498R and N501Y remained stable, likely contributing to stabilization of the BA.2.75 open trimer (Figure 3F). The RMSF profile for the S1 subunit (residues 14-365) in the BA.1 and BA.275 trimers featured fairly moderate values corresponding to residues 290-330. These residues include the N2R linker (residues 306-334) that connects the NTD and RBD within a single protomer. In the RBD-down conformation a segment of the N2R region forms a β strand (residues 311-318) that interacts with the CTD2 domain (residues 592-685) and β strand (residues 324-328) stacking against CTD1 (residues 528-591). These segments exhibit only moderate fluctuations in BA.1 (Figure 3B) and BA.2.75 open trimers (Figure 3F), while showing larger displacements in the BA.2 variant (Figure 3D). The central findings of the conformational dynamics analysis can be summarized as follows. First, the results showed the greater flexibility of the BA.2 trimer in both closed and open states. In contrast, we observed the increased stabilization of the BA.2.75 functional states as compared to the BA.1 and BA.2 structures. These results are in direct agreement with the recent functional studies, suggesting that BA.2.75 trimer is the most stable among these variants [50]. The dynamics of S Omicron BA.2 trimers showed a less compact inter-protomer packing as opposed to the tightly packed inter-subunit interfaces in BA.1 and BA.2.75 trimers. These dynamic characteristics gleaned from RMSF analysis are consistent with thermal stability assays that verified that S BA.2 trimers were the least stable among BA.1, BA.2, BA.3 and BA.4 variants [48]. The dynamics of BA.2.75 trimers is similar to BA.1 but is more stable as compared to BA.2 which is consistent with the experimental in vitro thermal stability data showing that the BA.2.75 S-trimer was the most stable among Omicron variants [50].

To identify hinge sites and characterize collective motions in the SARS-CoV-2 S Omicron structures, we performed principal component analysis (PCA) of trajectories derived from CG-BD simulations. The low-frequency modes can be exploited by mutations to modulate protein movements that are important for allosteric transformations between the open and closed states. The local minima along the slow mode profiles are typically aligned with the hinge centers, while the maxima could precisely point to the moving regions undergoing concerted movements. The functional dynamics in the S Omicron structures is highly conserved among variants and is illustrated for S BA.1 closed and open states (Figure 4). Interestingly, several Omicron sites belong to local hinge regions (N764K, D796Y, N856K, Q954H, N969K, and L981F), while RBD mutational sites are located in regions prone to functional movements. Additionally, and in line with our previous studies, we noticed that F318, S383, A570, I572 and F592, D985 residues are conserved hinge sites that are situated near the inter-domain SD1-S2 interfaces and could function as regulatory centers governing the inter-protomer and inter-domain transitions (Figure 4). Indeed, the disulfide bond between RBD and S2 at positions S383 and D985 stabilized the spike in an all RBD-down conformation. In the open forms of the S Omicron variants, we found that the RBD Omicron mutational sites occupy highly mobile regions that experience large displacements in collective motions, while mutational sites from the S2 subunit tend to reside in partly immobilized regions (Figure 4). Together, these results suggest a possible long-range coupling between stable S2 Omicron positions and highly mobile S1-RBD Omicron sites during conformational transitions. The coarse-grained nature of simulations and topology-centric picture of collective dynamics revealed a general trend shared across all Omicron subvariants. However, a more detailed energetic analysis of the inter-protomer interactions and changes induced by specific variants may introduce subtle changes in the stability and dynamic equilibrium between the open and closed states.

**Figure 4.**
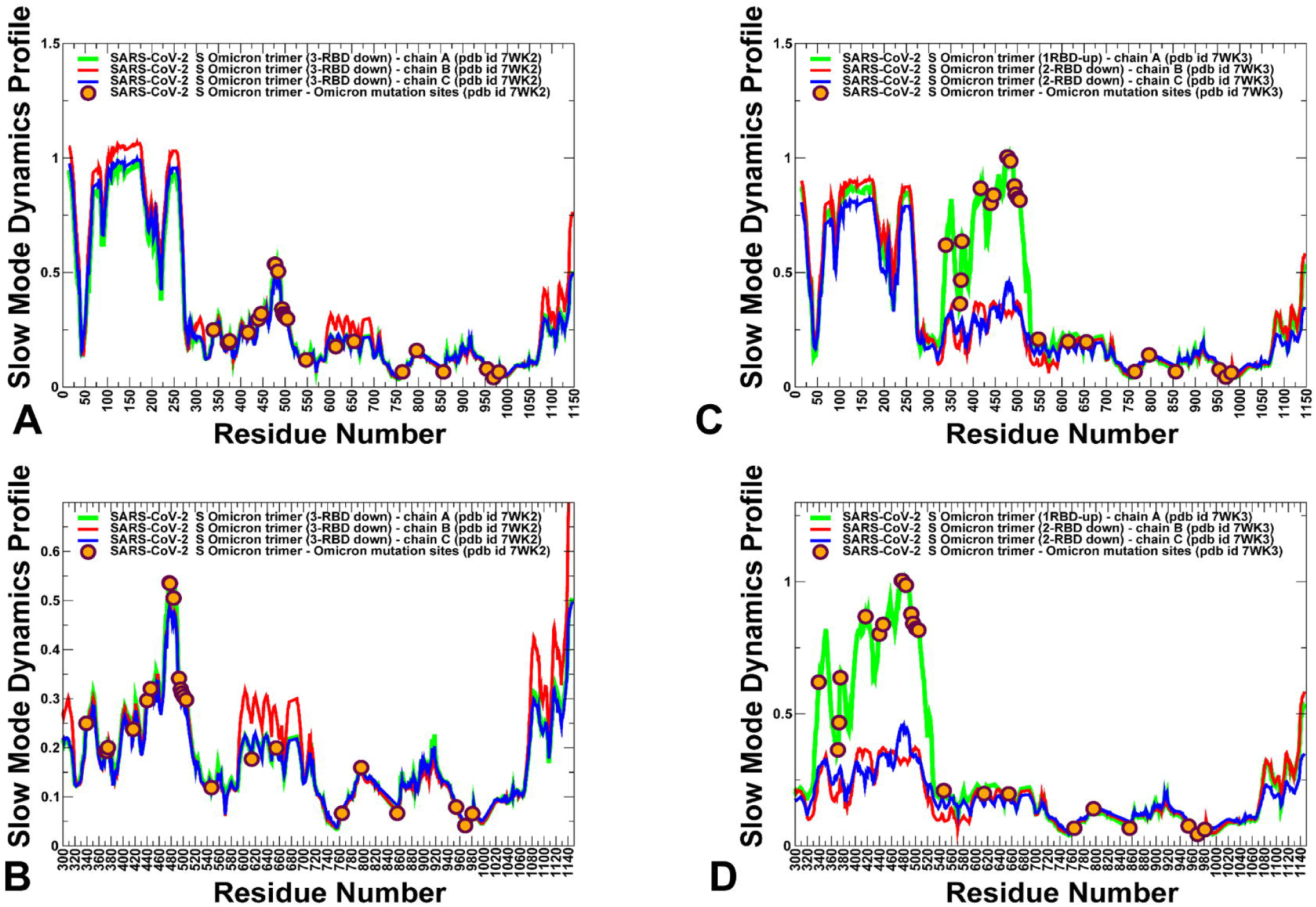
The slow mode mobility profiles of the SARS-CoV-2 S Omicron trimer structures. The slow mode dynamics profiles represent the displacements along slow mode eigenvectors and correspond to the of the slowest 10 modes. (A) The slow mode mobility profiles for the closed S Omicron BA.1 closed trimer structure (pdb id 7WK2). (B) A close-up of the slow mode profile for the RBD region in the BA.1 closed trimer. (C) The slow mode mobility profiles for the 1RBD-up open S Omicron BA.1 trimer structure (pdb id 7WK3). (D) A close-up of the slow mode profile for the RBD region in the BA.1 open trimer. The slow mode profiles for protomer chains A, B and C are shown in green, red and blue lines respectively. The positions of Omicron BA.1 mutational sites G339D, S371L, S373P, S375F, K417N, N440K, G446S, S477N, T478K, E484A, Q493R, G496S, Q498R, N501Y, Y505H, T547K, D614G, H655Y, N679K, P681H, N764K, D796Y, N856K, Q954H, N969K, and L981F are shown in orange-colored filled circles.

### Electrostatic Interactions in the Different Functional Forms of the SARS-CoV-2 Spike Omicron Trimers

To provide a detailed analysis of the electrostatic interactions in the S Omicron trimers, we employed the equilibrium ensembles obtained from CG-BD simulations. The electrostatic interaction potentials are computed for the averaged RBD-hACE2 conformations for by solving Poisson Boltzmann equation using the APBS-PDB2PQR software [86,87] based on the Adaptive Poisson–Boltzmann Solver (APBS) [86] and visualized using the VMD visualization tool [88]. Using this approach, we computed the electrostatic potential on the surface of the cryo-EM SARS-CoV-2 S trimer structures for the closed and open forms of the original strain Wu-Hu-1 (Figure 5A,B) and compared the distributions for the Omicron BA.1 closed and open states (Figure 5C,D), Omicron BA.2 variant (Figure 5E,F) and Omicron BA.2.75 states (Figure 5G,H). In agreement with some previous studies of the electrostatic potential on S proteins [75], we confirmed a significant change in the electrostatic distributions between Wu-Hu-1 trimers and S Omicron variants (Figure 5). The S Wu-Hu-1 trimer displayed mostly negatively charged electrostatic surface in the S2 and S1/S2 regions and only weak positively charged potential in the NTD and RBD regions (Figure 5A,B). In contrast, closed and open forms of the S Omicron BA.1 trimer showed a positively charged potential in the S1 subdomain, which becomes particularly strong in the S-NTD and S-RBD regions (Figure 5C,D). The Omicron BA.1 variant gained positive electrostatic potential from N440K, T478K and three novel mutations Q493R, Q498R and Y505H. A comparison of the electrostatic potential surfaces for the S Omicron variants showed a radical change in the distribution between Wu-Hu-1 and Omicron BA.1 but a more gradual evolution of the electrostatic potential between BA.1, BA.2 and BA.2.75. The overall differences in the electrostatic potential surfaces for the Omicron subvariants are relatively moderate, but we observed a greater accumulation of the positively charged regions in the S BA.1 trimers as compared to S BA.2 and S BA.2.75 trimers. The S BA.2.75 forms are characterized by more positively charged S1 regions (Figure 5G,H) as compared to somewhat weaker positive electrostatic potential in S BA.2 trimers (Figure 5E,F). In general, we found that the RBD binding interface is more positively charged in S BA.1 than in BA.2 and BA.2.75 trimers (Figure 5). Consistent with recent analysis [89–91], we found that the electrostatic potential in the S1 and S2 subunits and at the RBD the interface is more positive in BA.2 than in BA.2.75 but is less positive than in S BA.1 (Figure 5).

**Figure 5.**
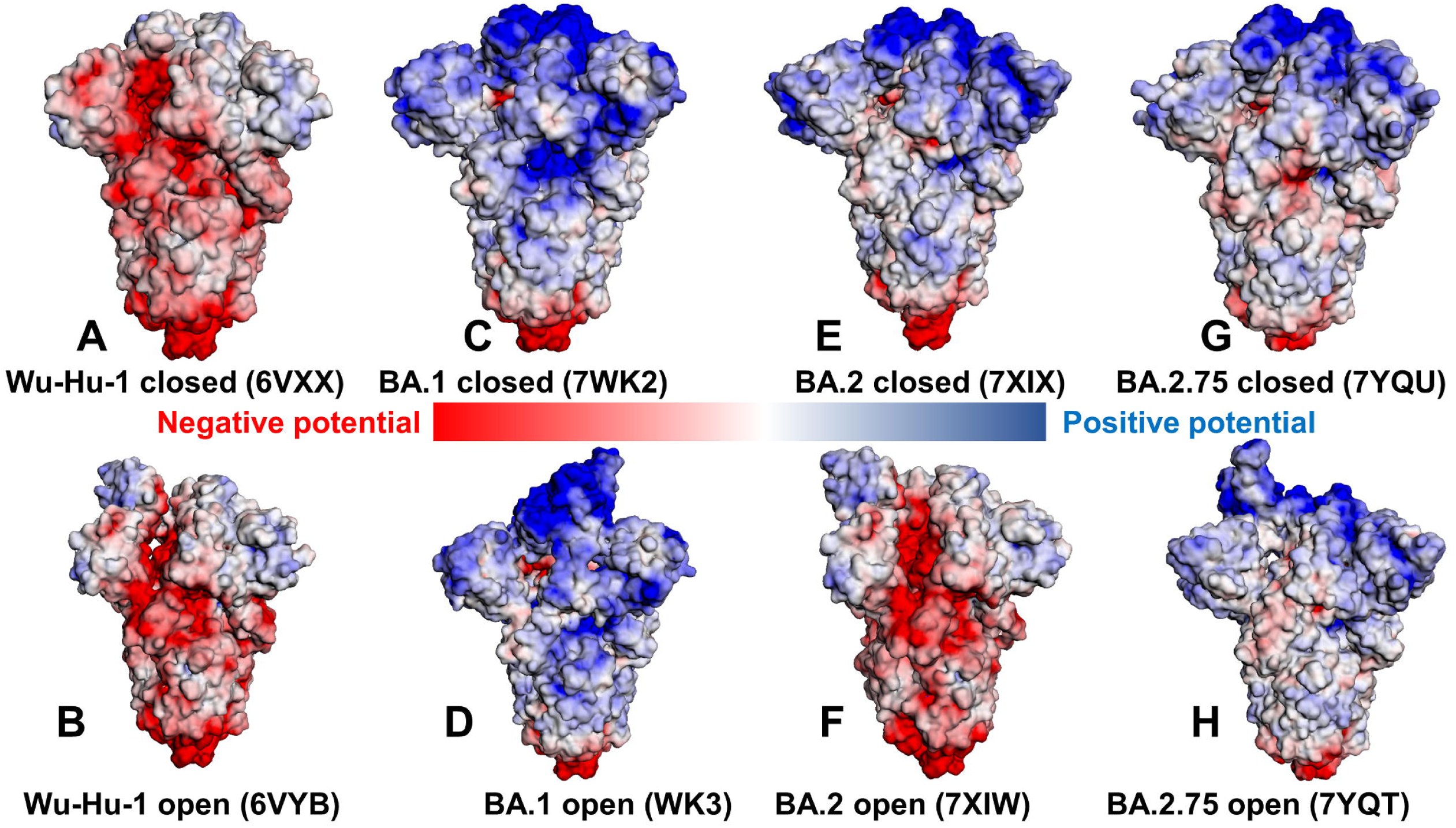
The distribution of the electrostatic potentials calculated with the APBS method [84,85] on the molecular surface of trimeric S proteins of Wuhan strain (Wu-Hu-1) closed state (pdb id 6VXX) (A), S-Wu-Hu-1 trimer in the open state (pdb id 6VYB)(B), S-Omicron BA.1 closed trimer (pdb id 7WK2) (C), S-Omicron BA.1 open 1-up trimer (pdb id 7WK3) (D), S-Omicron BA.2 closed trimer (pdb id 7XIX) (E), S-Omicron BA.2 open trimer (pdb id 7XIW) (F), S-Omicron BA.2.75 closed trimer (pdb id 7YQU) (G), S-Omicron BA.2.75 open trimer (pdb id 7YQT)(H). The color scale of the electrostatic potential surface is in units of kT/e at T = 37°C. Electro-positively and electronegatively charged areas are colored in blue and red, respectively. Neutral residues are white.

According to our analysis, the local increase of the positive electrostatic potential is partly caused by the presence of the positively charged Q493R in BA.1 and BA.2 that in BA.2.75 is reversed to the neutral Q493. BA.5 also conserves Q493 while Q493R mutation is present in BA.1, BA.2 and BA.3. Structural analysis showed that Q493R may form salt bridges with E35 on ACE2 that are absent in BA.2.75 due to reversed R493Q substitutions. Calculating the electrostatic potential, we also noticed that K460 of BA.2.75 S RBD is positively charged which may locally increase the positive electrostatic potential in this region, but this is offset by R493Q mutation. These structural observations suggest that while N460K substitution contributes to increased electrostatic complementary binding between the BA.2.75 S RBD and ACE2, this may be counterbalanced by the attenuated electrostatic interactions mediated by Q493. We also found that that S-BA.1 and S-BA.2 trimers displayed a greater net positive charge than S-BA.2.75 across the entire S protein, including S1 and S2 regions, suggesting that electrostatic differences are distributed through the entire S protein beyond the RBD binding interface with the ACE2 receptor (Figure 5).

In general, despite mutational differences between BA.1, BA.2 and BA.2.75 lineages, the electrostatic potential surfaces for the respective S trimer structures showed a strong positive electrostatic surface in the S-RBD regions which interface with the negatively charged ACE2 receptor. The results are generally consistent with the hypothesis about the relationship between the enhanced electropositive character of the RBD variants and their enhanced capacity for the enhanced transmission [92]. Despite changes in the RBDs and in other regions of the S protein in the Omicron BA.1, BA.2 and BA.2.75 trimers, the net charges for these lineages remain relatively similar and the molecular electrostatic surfaces of S homotrimers for BA.1, BA.2 and BA.2.75 lineages are only marginally different. According to the results, the enhanced electropositive character of the RBDs due to single or only several critical mutations may be sufficient to infer the improved binding towards the strongly electronegatively charged ACE2 receptor (Figure 5). Our analysis also suggested that the improved binding affinity of the BA.2.75 variant with ACE2 may not be solely determined by the electrostatic changes as N460K and R493Q may act in opposite directions in modulating the positive electrostatic potential on the RBD. We argue that evolution of the SARS-CoV-2 virus changes for newly emerging variants may have reached a certain critical plateau of electrostatic positively charged RBD distribution that is optimal to complement ACE2 electrostatic potential, and convergent mutations could have emerged to balance multiple tradeoffs rather than progressively improving the RBD-ACE2 binding affinities.

### Mutational Scanning and Sensitivity Analysis Identify Key Structural Stability Hotspots in SARS-CoV-2 S Omicron Subvariants

To dissect the energetics and determine the stability profiles of the S trimers for the Omicron variants, we employed two different approaches with the increasing level of complexity and rigor. In silico mutational scanning of the S trimer residues was first done using knowledge-based BeAtMuSiC approach [93–95]. This approach allows for fairly robust and accurate predictions of the effect of mutations on both the strength of the binding interactions and on the stability of the complex using statistical potentials and neural networks. The adapted in our study approach was enhanced by applying the ensemble-based averaging of free energy computations using 1,000 samples from simulation trajectories. Here, we focused on fast and systematic mutational scanning and sensitivity analysis for the inter-protomer interface residues (Figure 6) as the resulting profiles can illustrate the density of the inter-protomer contacts in different regions and single out specific inter-protomer hotspots in which mutations on average cause significant energetic perturbations. The results of this analysis are presented in the form of scatter plots for the open and closed trimers of the S Omicron BA.1 (Figure 6A,B), BA.2 (Figure 6C,D) and BA.2.75 variants (Figure 6E,F). First, we found that mutation-sensitive positions in the S trimers were distributed across both S1 and S2 subunits even though S2 is a considerably more rigid subunit as compared to a more dynamic S1 subunit (Figure 6). Second, for all S trimers, there is a clear density of potential stability hotspots localized in the S1-RBD core regions and at the RBD inter-protomer regions that are often structurally proximal to highly dynamic RBM regions at the top of S1 subunit. Interestingly, it could be seen that the distributions of stability centers in the NTD and RBD regions become denser in the open states of the S trimers, owing to the RBD-RBD inter-protomer contacts (Figure 6). Indeed, mutations of the NTD residues are generally tolerant in the closed trimers where NTDs are not involved in the inter-protomer packing but become somewhat more sensitive in the open states where RBD-up conformation is involved in the intra-molecular contacts. However, the free energy changes caused by mutations of the NTD residues remain fairly moderate with the exception of F43, Y200 and F201 (Figure 6). The RBD core α-helical segments (residues 349-353, 405-410, and 416-423) formed a consistent cluster of energetic hotspots in all S trimers. The stability of the central β strands (residues 354-363, 389-405, and 423-436) is also evident from the folding free energy changes, indicating that the integrity of the RBD core is preserved in both open and closed states for all Omicron variants (Figure 6). Other conserved stability regions shared among functional states of the S trimers for all Omicron variants include clusters in the CTD1 (residues 535-560), UH (residues 700-780) and CH regions (residues 986-1000). Several positions that are particularly sensitive to mutations leading to large destabilization changes included highly conserved F377, Y421, F592, Y707, and Y917 (Figure 6). The S trimer residue F43 lies at an interface between adjacent monomers and plays an important role in modulating inter-protomer packing and antibody binding. In fact, the recent experimental studies that employed alanine-scanning loss-of-binding experiments for COV2-3434 and COV2-3439 antibodies identified the primary residues F43, F175, L176, and L226 as critical for binding of COV2-3434 [96]. Despite the presence of commonly shared stability clusters, the results revealed denser distributions for the BA.1 trimers (Figure 6A,B) and BA.2.75 trimers (Figure 6E,F).

**Figure 6.**
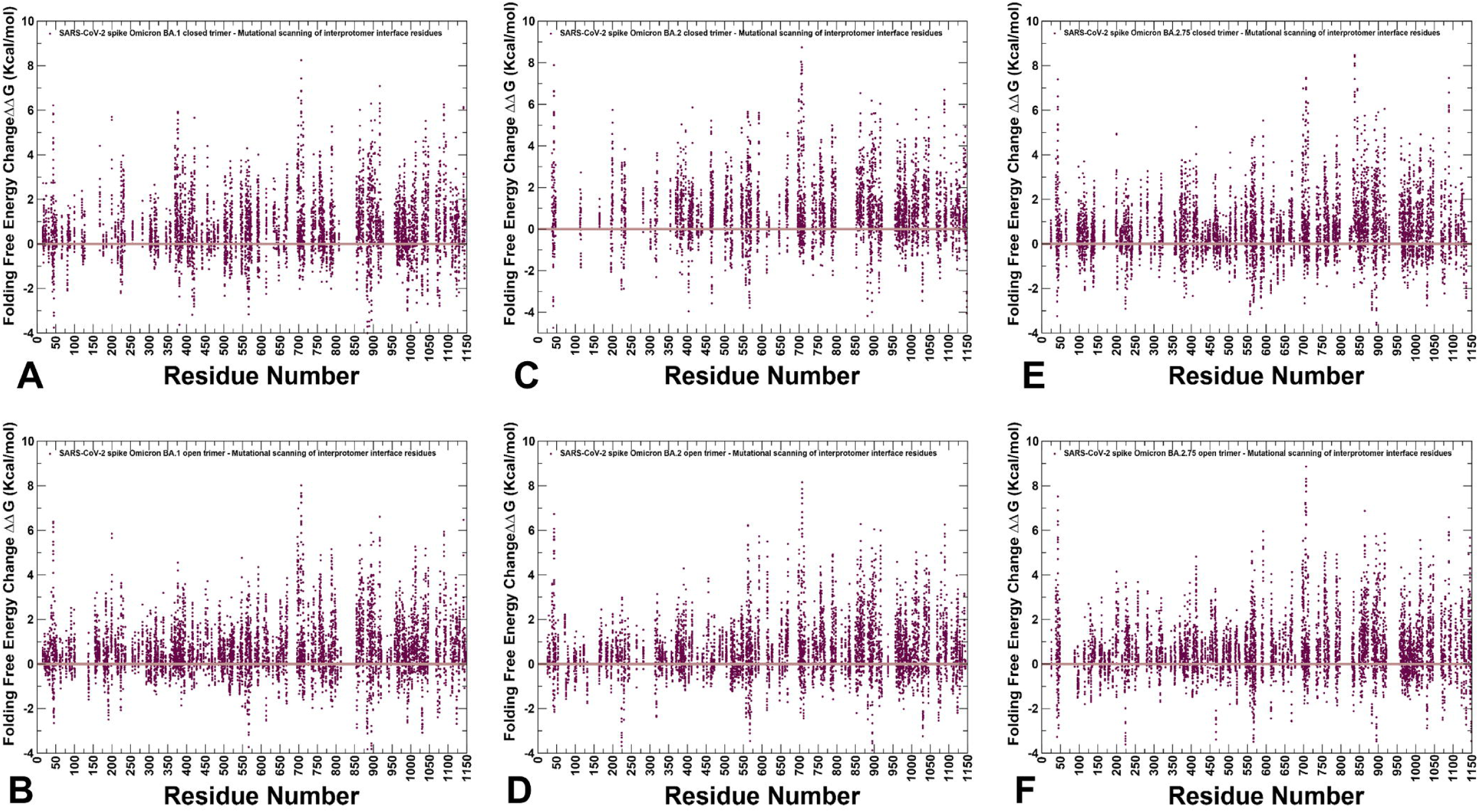
Ensemble-based mutational scanning of protein stability and binding for the SARS-CoV-2 S Omicron trimers. The mutational scanning scatter plot for the SARS-CoV-2 Omicron BA.1 closed trimer, pdb id 7WK2 (A), for the S Omicron BA.1 trimer in the open form, pdb id 7WK3 (B), for the SARS-CoV-2 S Omicron BA.2 closed trimer, pdb id 7XIX (C), for the S Omicron BA.2 trimer in the open form, pdb id 7XIW (D), for the SARS-CoV-2 S Omicron BA.2.75 closed trimer, pdb id 7YQU (E), for the S Omicron BA.2.75 trimer in the open form, pdb id 7YQT (F). The standard errors of the mean for binding free energy changes were based on a different number of selected samples from a given trajectory (500, 1,000 and 2,000 samples) are within 0.12-0.18 kcal/mol.

Importantly, the results of mutational scanning confirmed that the closed and open forms of the BA.2 variant are more dynamic as the S-BA.2 residues appeared to be more tolerant to substitutions (Figure 6C,D). Interestingly, the folding free energy distribution is markedly sparser in the closed trimer for BA.2 trimer (Figure 6C), suggesting that the density of the inter-protomer contacts is weaker reflecting the increased mobility of this BA.2 form. This may imply that the closed form is less stable, and the equilibrium can be more easily shifted to the 1RBD-up open state. The distribution of the inter-protomer energetic hotspots in the open BA.2 form is comparable to the other Omicron variants but showed a weaker density in the NTD and RBD regions of the S1 subunit (Figure 6D). These results imply that the S BA.2 variant states may be less stable as compared to the S BA.1 and BA.2.75 trimers, which is consistent with the emerging experimental evidence on thermostability of the Omicron variants [48–50].

By employing the conformational ensembles of the S trimers, we also performed systematic mutational scanning of the interprotomer residues and computed folding free energy changes using FoldX approach [97–100]. The adapted in our study modification of FoldX approach is enhanced through ensemble-based averaging of the folding free energy computations. We report the complete “deep” scanning energy heatmaps that include a full list of the S residues for each variant that are involved in the inter-protomer contacts (Supporting Information, Figures S1-S6). To focus on a sensible number of important for protein stability residues, these complete heatmaps represent systematic mutational scanning for residues in which mutations produced significant destabilization changes in the range of 2.5 kcal/mol < ΔΔG values < 9 kcal/mol. To make these heatmaps more informative, the free energy changes that are outside of this range are highlighted in white color. This allows us to dissect the main trends in the mutational scanning of protein stability and characterize groups of important stability hotspots (Supporting Information, Figures S1-S6). To facilitate a comparison and highlight the most dominant centers of protein stability, we also reported the mutational scanning heatmaps that are deliberately filtered to focus on residues where mutations induce extremely large destabilization changes with ΔΔG values > 5.0 kcal/mol (blue scale) as well as positions and mutations which yielded significant stabilization changes ΔΔG values < −3kcal/mol (yellow scale) (Figure 7). In order to highlight key positions and respective modifications leading to these effects, positions/mutations that do not meet these thresholds are shown in white.

**Figure 7.**
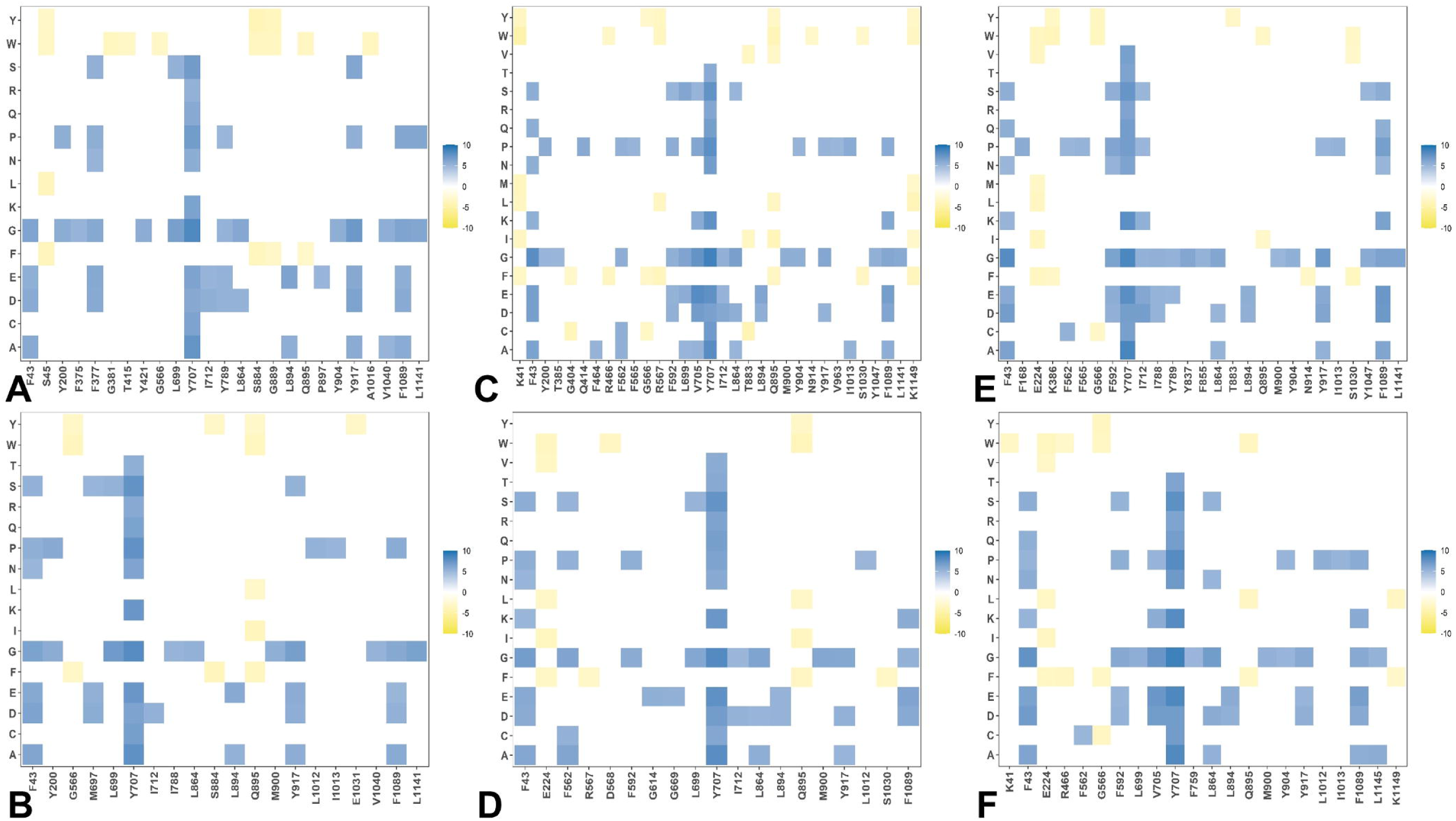
Ensemble-based dynamic mutational profiling of the S trimer inter-protomer interfacial interfaces in the S Omicron trimers. The mutational scanning heatmaps are shown for the inter-protomer residues in the S Omicron BA.1 closed trimer, pdb id 7WK2 (A), S Omicron BA.1 open trimer, pdb id 7WK3 (B), S Omicron BA.2 closed trimer, pdb id 7XIX (C), S Omicron BA.2 open trimer, pdb id 7XIW (D), S Omicron BA.2.75 closed trimer, pdb id 7YQU (C), S Omicron BA.2.75 open trimer, pdb id 7YQT (D). The standard errors of the mean for binding free energy changes were based on MD trajectories and selected samples (a total of 1,000 samples) are within ∼ 0.15-0.23 kcal/mol using averages over 10 different CG-BD trajectories. The mutational scanning heatmaps are filtered to focus on residues where mutations induce extremely large destabilization changes with ΔΔG values > 5.0 kcal/mol (blue scale) as well as positions and mutations which yielded significant stabilization changes ΔΔG values< −3kcal/mol (yellow scale). The positions/mutations that do not meet these thresholds are shown in white. According to the maps, only about 30 residues from all structures were identified as important.

The first important observation of this analysis was the presence of denser network of inter-protomer interactions in the open and closed forms of BA.1 (Figure 7A,B, Supporting Information Figure S1,S2) and BA.2.75 trimers (Figure 7E,F, Supporting Information Figure S5,S6). To streamline the analysis of the inter-protomer interfaces, we highlighted the S residues in which mutations may induce significant destabilization and result in the large positive free energy changes > 2.5 kcal/mol (Supporting Information, Figures S1-S6). The hotspot clusters are broadly distributed, including the RBD residues and CTD1 domain (residues 529-591) but most of the hotspots are localized in the S2 subunit regions, including the HR1 (residues 910-985), CH regions (residues 986-1035), and UH regions (residues 736-781). Of particular interest is the existence of a shared cluster of stability hotspots that includes M697, L699, G700, V705, Y707, and I712 (Supporting Information, Figures S1-S6). The folding stability maps for the BA.1 closed trimer revealed a number of significant stability centers, particularly F43, Y369, F377, Y421, F592, L699, Y707, I712, G757, Y789, W886, Y917, L1034 and F1089 (Supporting Information, Figures S1, Figure 7A). Mutations of these residues often produce large destabilization effect, and the respective sites represent sensitive energetic hotspots. Some of these hotspot centers are shared in the open BA.1 trimer including F43, F377, F592, L699, Y707, G757, Y917, and F1089 (Supporting Information, Figure S2 and Figure 7B). In addition, we found that mutations in the RBD sites F456, N487 and Y489 may also lead to large destabilization effects. Importantly, protein stability hotspots in the BA.1 trimers included Y789, F592, F562, V382, S383, and D571 positions that correspond to functionally important hinge regions. Strikingly, one of these vital sites includes global hinge F592 position that is strategically positioned at the inter-domain interfaces and interacts with N856, G857, K854, F855 and M740 residues that collectively form an important regulatory cluster of the inter-domain movements controlling functional changes between the open and closed states. Another interesting observation is that sites of Omicron mutations do not typically correspond to the major stability hotspots, but some of the Omicron positions are involved in direct contacts with the hotspots. The Omicron mutational sites are involved in the inter-protomer bridges formed near stabilization hotspots and include N764K-Q314, S982-T547K, N856K-D568, N856K-T572, N969K-Q755, N969K-Y756, S383-D985, F855-G614, V963-A570, N317-D737, R319-D737, R319-T739, R319-D745, and K386-F981. These inter-protomer bridges are anchored by the Omicron sites (T547K, D614G, N764K, N856K, N969K, and L981F) from the S2 subunit. However, our results also highlighted a noticeably sparser distribution of large destabilization changes in the BA.2 states (Supporting Information, Figures S3,S4) which is reflective of the increased flexibility of the BA.2 trimer. A close inspection of these maps for the BA.2 trimer (Supporting Information, Figures S3,S4) indicated that fewer residues can be assigned to potential hotspots and many sensitive positions, particularly in the RBD core and S2 regions become more tolerant. This becomes especially apparent in the open form of BA.2 trimer (Supporting Information, Figure S4 and Figure 7D), thus supporting the notion that the decreased thermal stability ty of this variant may be linked with the greater mobility and significant conformational heterogeneity of the open state [48–50]. It is possible that these dynamic and energetic signatures may facilitate acquisition of the ACE2-exposed open form and lead to the increased infectivity exemplified by BA.2. Notably, however, the key hotpots are preserved including sites associated with the inter-protomer interactions and hinge regions (F318 and F592). The heatmap for the closed BA.2.75 trimer (Supporting Information, Figures S5, Figure 7E) is similar to that of the BA.1 and BA.2 trimers. Nonetheless, we noticed some relevant differences in the S2 regions, particularly the increased density for residues F592, M697, L699, G700, V705, Y707, and I712 (Supporting Information, Figure S5). More important changes were observed for the open S BA.2.75 trimer (Supporting Information, Figure S6), where we observed significant energetic destabilization caused by mutations of the NTD positions (N125,Y160,G199,Y200, Y204, P225, V227, L229,G257,G283) as well as some key RBD residues (F318, R319, V382, L390, Y396, F464). Overall, we found the increased number of positions in the open BA.2.275 trimer that are mutation-sensitive, indicating that BA.2.75 trimer may be more stable than BA.2 and comparable in stability to BA.1 variant.

Of notice is the observation that Q613 and D614G positions correspond to strong energetic hotspots in the open states of all Omicron variants, showing that D614G modification can significantly contribute to the stability of the 1RBD-up functional spike state. Structurally, the D614G mutation may weaken some interprotomer contact, which caused the RBD to more readily adopt RBD-up conformation, thus facilitating the interactions with the ACE2 receptor. However, D614G plays role in stabilizing the noncovalent association of the S1 and S2 spike subunits. Our data agree with the experimental data showing that D614G mutation makes the S protein more stable but also more prone to interact with its receptor [101]. Interestingly, we found that mutations of serine RBD sites S371, S373 and S375 (S371L, S373P and S375F in BA.1; S371F, S373P, S375F in BA.2; S371F, S373P, S375F in BA.2.75) represent moderate energetic hotspots in the closed forms, while in the open states these positions are more tolerant (Supporting Information, Figures S1-S6). This is consistent with structural analyses showing that mutations of S371L, S373P, and S375F can promote the interprotomer interactions between the down RBDs and closer packing of the RBD-RBD interface via interactions of S373P and S375F with the N501Y and Y505H substitutions in the adjacent RBD [27,31]. A systematic functional analysis of Omicron subvariants BA.1 and BA.2 on S protein infectivity and neutralization showed that individual mutations of S371F/L, S375F, and T376A as well as S2 mutations Q954H and N969K in impaired infectivity, while changes to G339D, D614G, N764K, and L981F can moderately enhance it [102].

Instead of targeting sites that are highly sensitive to mutations affecting the global protein stability, Omicron mutations often emerge in structurally-proximal to the hotspots positions which allows for modulation of stability and tradeoffs between stability, adaptability and antibody binding. Of particular importance is the role of the most dominant stability hotspot Y707 that is shared across all variants (Figure 7, Supporting Information, Figures S1-S6). Y707 forms a number of important direct inter-protomer contacts including T883, I896, Y796 and other residues. Strikingly, Y707 forms an important inter-domain bridge Y707-D796 in the native spike that becomes partly disrupted upon D796H/Y mutations. Moreover, D796H/Y changes that disrupt an inter-domain D796-Y707 bridge can act allosterically in conferring virus resistance to neutralization [103]. It was reported that mutant D796H may be an important contributor to the decreased susceptibility to neutralizing antibodies but may lead to reduced infectivity [104]. According to our results, D796Y Omicron mutation shared among BA.1, BA.2 and BA.2.75 can alter the native contacts with the central stability hub Y707 and through these interactions allosterically modulate balance of dynamics, stability and long-range communications in the S2 subunit. Other S2 Omicron mutations including N856K, Q954H and N969K may operate similarly, where instead of directly targeting sensitive stability hotspots, these modifications can alter interactions mediated by the major energetic hubs (F592, Y707, Y917). In particular, one of the major stability hubs F592 belongs to the inter-domain S1-S2 hinge cluster and is involved in a number of inter-protomer clusters (B855-C589 C592; A740-A856-B592; A592-C737-C855). Through these stable clusters F592 interacts with N856 (and N856K in BA1, BA.2 and BA.275). N856K Omicron mutation may alter local interactions and stability of clusters anchored by F592 and as a result perturb global conformational changes and long-range signaling controlled by F592 site. Our results suggest that Omicron sites are coupled to major stability hotspots through local interactions may allow for modulation of global stability and dynamic changes in the S protein without significantly compromising the integrity and activity of the spike. These results further highlight the underlying mechanism of S proteins that is linked with the subtle balancing and tradeoff between protein stability, conformational adaptability and plasticity as well as immune escape pathways that collectively drive evolution of the SARS-CoV-2 S variants and emergence of BA.2 and BA.2.75 subvariants that are markedly different in their effect on dynamics and stability.

While our analysis cannot directly address functional roles of specific Omicron mutations in complex functional responses, we found that a delicate balance of moderate thermodynamic stability and sufficient degree of conformational adaptability is characteristic of most Omicron mutational sites which may allow these positions to modulate protein responses through couplings with major stability centers. To explore a hypothesis that Omicron sites may be coupled with the stability centers in a functionally relevant dynamic network, we mapped the Omicron BA.1 mutational sites and protein stability hotspots on structures of the closed BA.1 trimer (Figure 8A,B) and open BA.1 trimer (Figure 9A,B) it can be seen that these important functional sites have only partially overlap but may play complementary roles in modulating spike functions and binding. An important revelation of this analysis was that structural stability hotspots can be interconnected and form potential communication routes between functional regions of the S2 and S1 subunits (Figures 8,9). Moreover, it appeared that structural stability centers also link the inter-protomer and inter-domain regions along fairly narrow “communication pathways” in the closed Omicron trimer (Figure 8B). A similar functional topography of stability centers could be seen in the open Omicron trimer (Figure 9B) where these sites link the S1 and S2 regions along potential communication spine of the S protein. The results suggested that stability hotspots may be important not only for structural integrity and thermostability of S proteins but could also form “stepping stones” for mediating long-range allosteric communications between S1 and S2 subunits.

**Figure 8.**
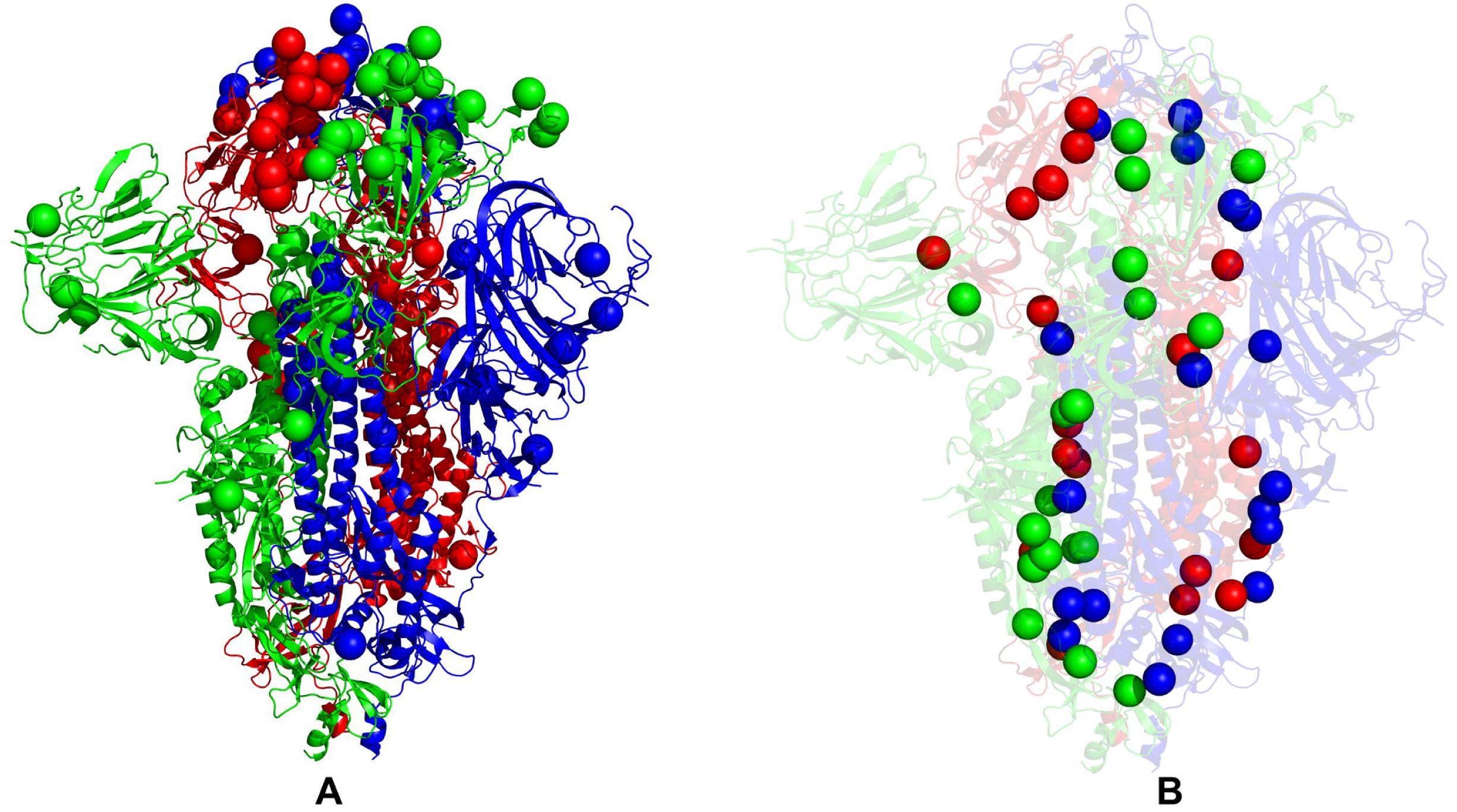
Structural maps of the functional sites for the SARS-CoV-2 S Omicron BA.1 closed trimer. (A) The structural mapping of the Omicron sites projected onto the cryo-EM structure of the closed S Omicron trimer (pdb id 7WK2). The protomers are shown in green, blue and red ribbons. The Omicron mutational sites G339D, S371L, S373P, S375F, K417N, N440K, G446S, S477N, T478K, E484A, Q493R, G496S, Q498R, N501Y, Y505H, T547K, D614G, H655Y, N679K, P681H, N764K, D796Y, N856K, Q954H, N969K, and L981F are shown in spheres for all protomers and colored according to the protomer identity. (B) The structural mapping of the protein stability hotspots (Supporting Information, Figure S1) onto the cryo-EM structure of the closed S Omicron trimer (pdb id 7WK3). The stability hotpots are shown in spheres colored according to the protomer identity.

**Figure 9.**
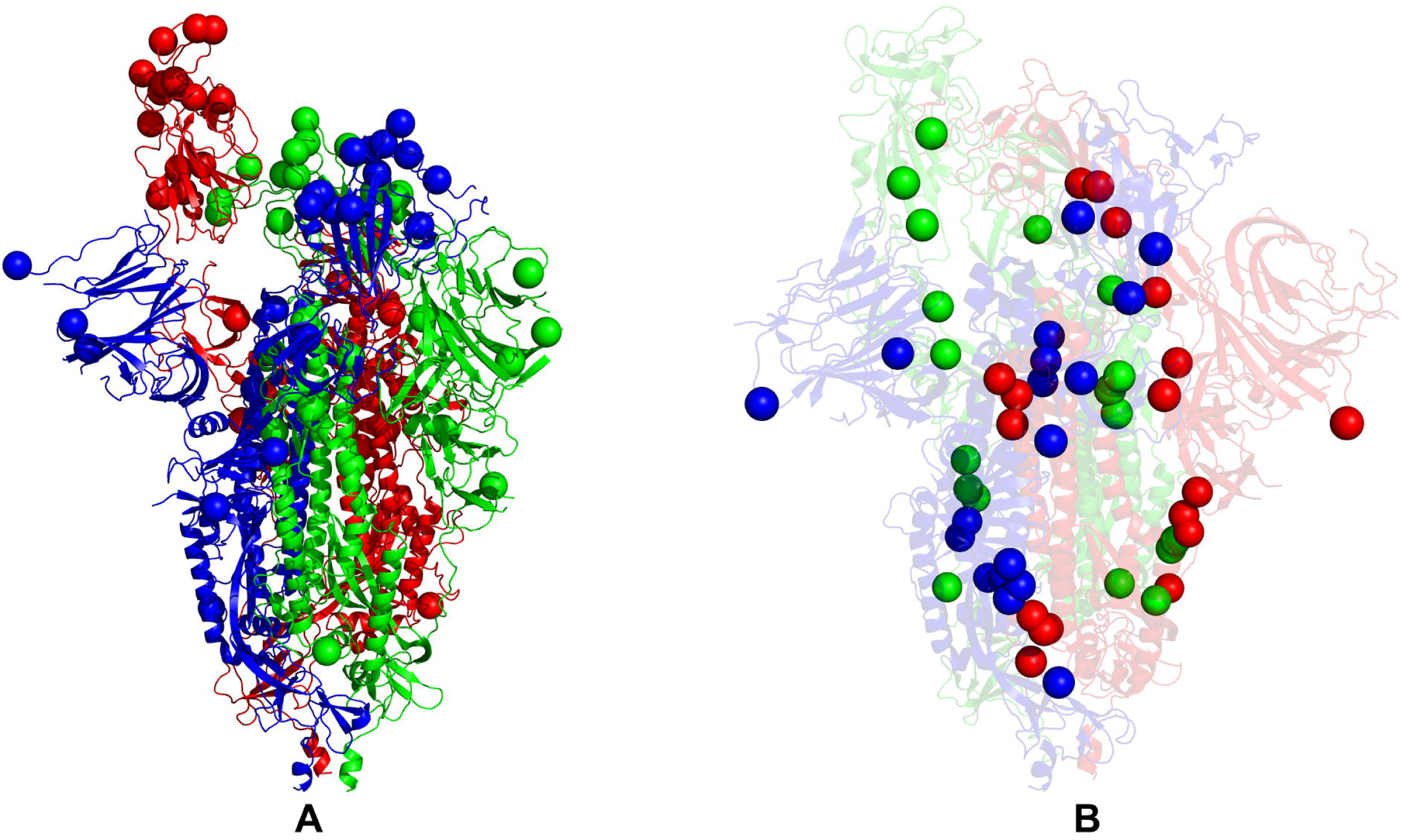
Structural maps of the functional sites for the SARS-CoV-2 S Omicron BA.1 open trimer. (A) The structural mapping of the Omicron sites projected onto the cryo-EM structure of the open S Omicron trimer (pdb id 7WK3). The protomers are shown in green, blue and red ribbons. The Omicron mutational sites G339D, S371L, S373P, S375F, K417N, N440K, G446S, S477N, T478K, E484A, Q493R, G496S, Q498R, N501Y, Y505H, T547K, D614G, H655Y, N679K, P681H, N764K, D796Y, N856K, Q954H, N969K, and L981F are shown in spheres for all protomers and colored according to the protomer identity. (B) The structural mapping of the protein stability hotspots for the open BA.1 trimer, pdb id 7WK3 (Supporting Information, Figure S2). The stability hotpots are shown in spheres colored according to the protomer identity.

The stability of these S residues and their ordered interconnectivity may ensure efficient and rapid signal transmission. At the same time, we noticed that structural distribution of the Omicron sites in the S2 regions where these sites are located in physical proximity of major stability centers may allow for modulation of long-range interactions and broadening of the ensemble of putative communication routes. While the presence of significant number of S1-RBD Omicron sites is mainly associated with the modulation of binding affinities with the ACE2 and antibodies, these flexible positions may be dynamically coupled to the “rigid” communication routes formed by the stability hotspots, leading to modulation of long-range dynamic responses of the S Omicron trimers.

Concurrently, we also employed simplified SWOTein predictor of stability which identifies the residue contributions to the global folding free energy through three types of database-derived statistical potentials that include inter-residue distances, backbone torsion angles and solvent accessibility [105]. According to this approach, positive folding free energy contributions indicate stability weaknesses while large negative folding free energy contributions for a given residue suggest stability strength for this position [105,106]. We applied this approach to calculate the folding free energy contributions of each residue in the open and closed conformations of the S trimers. This analysis is similar to our study of conformational and mutational frustration in S proteins in which positions of minimal frustration are typically associated with the stability centers, while more energetically frustrated residues point to more dynamic and tolerant to mutations sites [107]. Similarly, as in the ensemble-based mutational scanning, we enhanced SWOTein computations by considering ∼1,000 representative conformations from sampling trajectories and then taking the average values of the folding free energies (Figure 10). Despite its simplicity, this approach considers key contributions to the folding free energy associated with the enthalpic components, hydrophobic interactions and entropic estimations. The stability strengths and weaknesses are identified as residues that upon mutation result in strong destabilization or strong stabilization The results are generally consistent with the more detailed mutational scanning analysis and highlighted the density of stable residues (highly negative folding free energy values) broadly distributed in the S1 and S2 subunits (Figure 10). The clusters of strong stability centers include residues in the RBD core (residues 428-434), and residues 545-562 from CTD1. The analysis of weak and strong spots of protein stability revealed a similar set of major stability centers located in highly conserved F377, Y421, F592, Y707, and Y917 residues as well as strong stability hotspot at D614G site (Figure 10). We also observed a considerable amount of energetic frustration (i.e., weak stability sites with positive folding free energies) across all regions, particularly in the NTD regions. These revelations suggested that S protein dynamics and functions may be governed by a complex interplay between structurally stable and more flexible S regions that collectively determine functional changes, balancing the requirements for stability, adaptability and binding.

**Figure 10.**
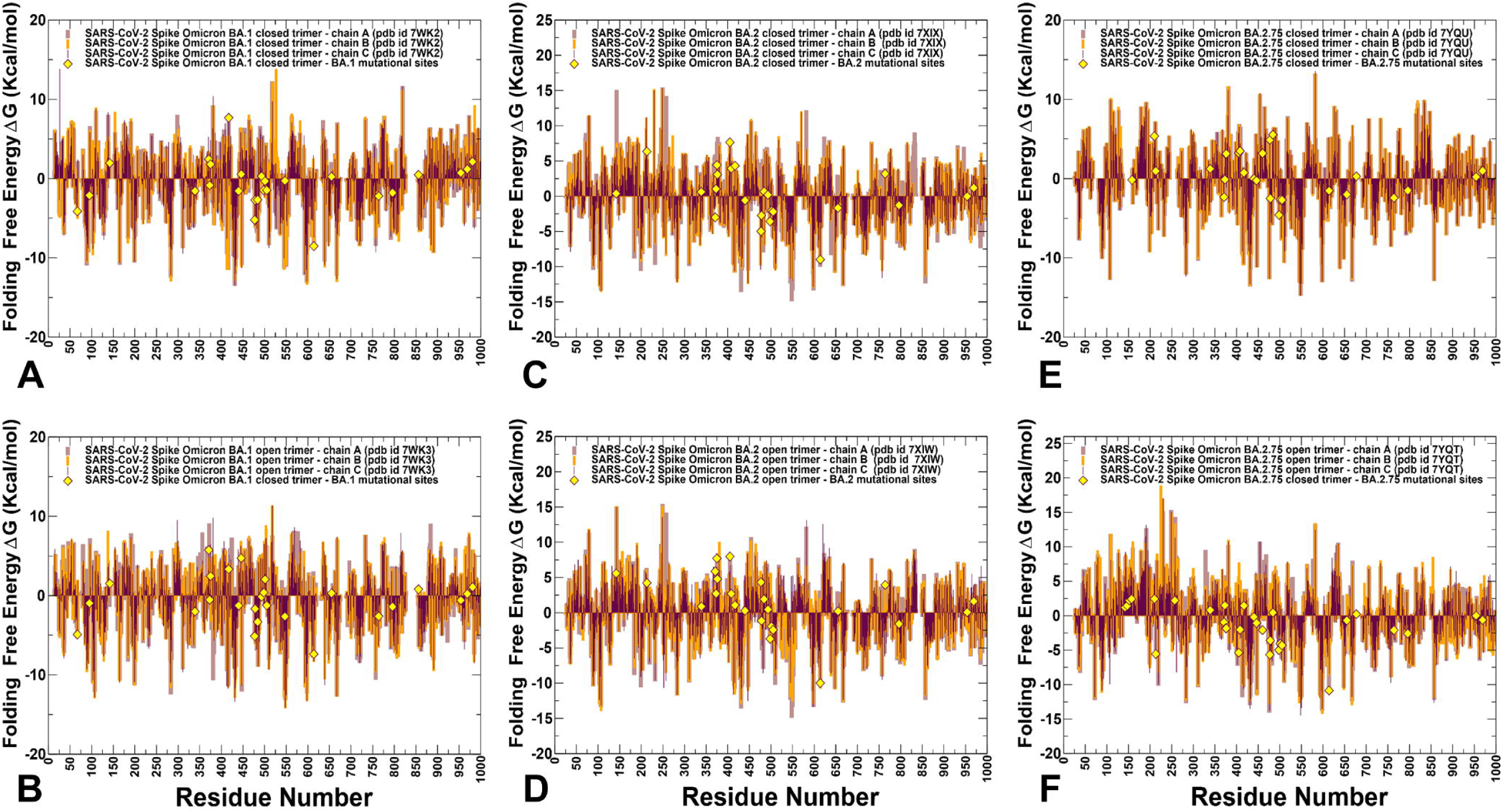
The folding free energies of the S protein residues for the SARS-CoV-2 S Omicron BA.1, BA.2 and BA.2.75 trimers. The folding free energies for S BA.1 closed trimer, pdb id 7WK2 (A), S BA.1 open trimer, pdb id 7WK3 (B), S BA.2 closed trimer, pdb id 7XIX (C), S BA.2 open trimer, pdb id 7XIW (D), S BA.2.75 closed trimer, pdb id 7YQU (E) and S BA.2.75 open trimer, pdb id 7YQT (F). The profiles are shown in brown, maroon and orange-colored bars for protomers A, B, and C respectively. The positions of Omicron mutational sites on the profiles are highlighted in yellow-colored filled diamonds. The projection of Omicron sites is shown for protomer A.

Notably, the analysis showed that highly stable and flexible positions in the S trimers intertwine and are often in local proximity of both sequence and structural spaces. Somewhat surprisingly, in a generally stable S2 subunit, there are also a significant number of weak centers with a considerable degree of frustration and intrinsic mobility (Figure 10). It is worth noting that due to the simplicity of this approach, it captures the main patterns of stability and flexibility, but may be less sensitive to subtle changes incurred by the Omicron variants. Nonetheless, by mapping sites of Omicron mutations for BA.1, BA.2 and BA.2.75 on the distributions, we found that the majority of these sites correspond to moderately weak centers, implying that mutations in these positions may improve stability of the S protein. It could be also noticed that in the BA.2.75 open and closed states, a number of Omicron mutational sites in S1 and S2 subunits featured moderately negative folding free energies, which reflected the marginally increased stability of these residues in BA.2.75 (Figure 10). These findings are consistent with the mutational scanning analysis and in agreement with the experimental evidence showing the increased folding stability of the S Omicron BA.2.75 trimers.

### Dynamic Network Modeling and Short Path Centrality Analysis Identify Regulatory Regions Mediating Allosteric Interaction Networks in the SARS-CoV-2 Spike Mutants

The residue interaction networks in the SARS-CoV-2 spike trimer structures were built using a graph-based representation of protein structures [108,109] in which residue nodes are interconnected through both dynamic [110] and coevolutionary correlations [111]. The dynamic residue interaction networks are constructed using an approach in which the dynamic fluctuations obtained from simulations are mapped onto a graph with nodes representing residues and edges representing weights of the measured dynamic properties. By employing perturbation-based network modeling, we computed ensemble-averaged distributions of the short path betweenness residue centrality (Figure 11). This network metric was explored to identify mediating centers of allosteric interactions in the SARS-CoV-2 Omicron BA.1, BA.2 and BA.275 structures. By using network modeling, we characterized the organization of residue interaction networks in the S Omicron functional states. We examined a possible role of Omicron mutations in allosteric interaction networks, based on a hypothesis that Omicron variants may induce control on a balance and trade-offs between conformational plasticity and protein stability.

**Figure 11.**
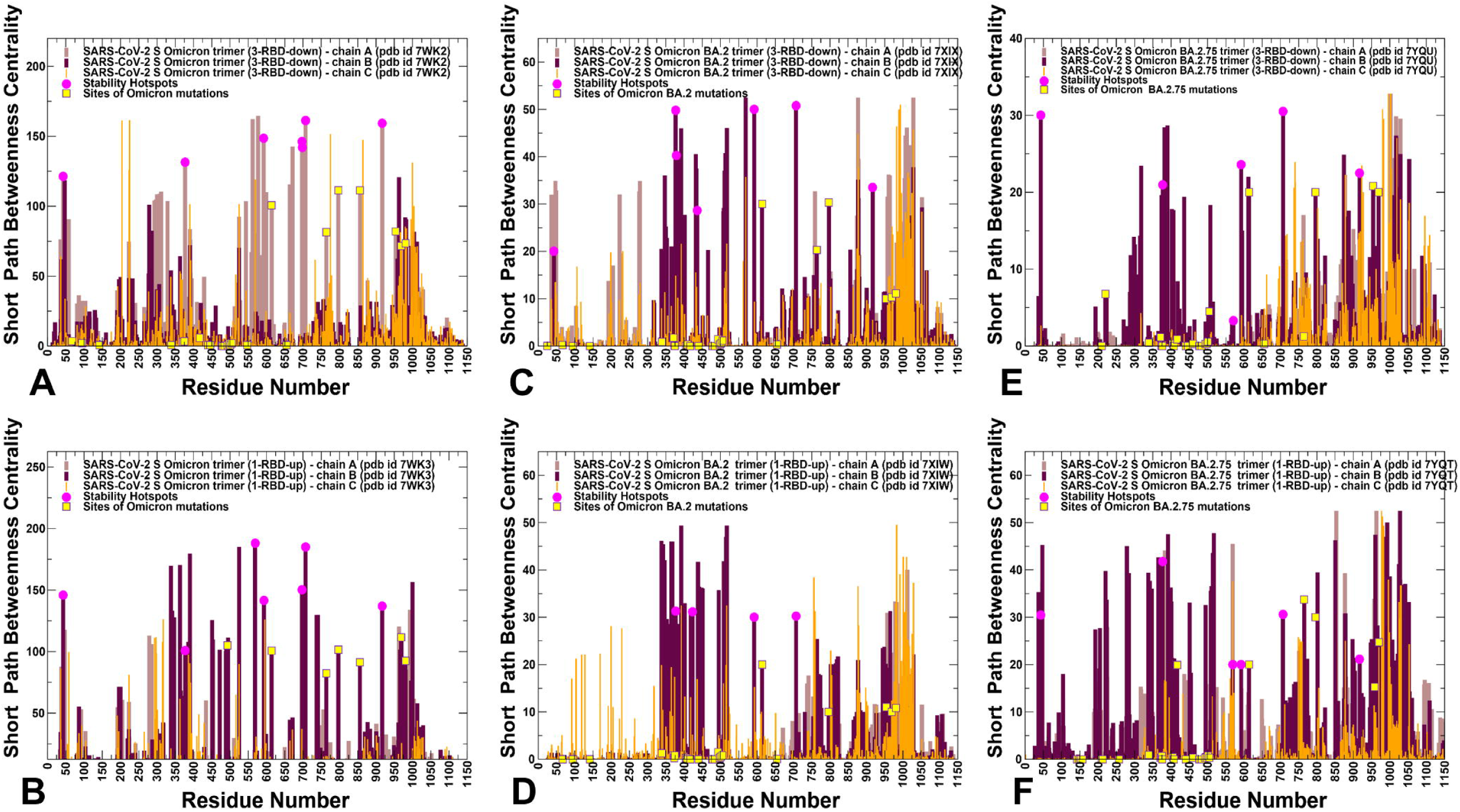
The short path betweenness centrality distributions for the SARS-CoV-2 S Omicron structures. The centrality profiles for the for S BA.1 closed trimer, pdb id 7WK2 (A), S BA.1 open trimer, pdb id 7WK3 (B), S BA.2 closed trimer, pdb id 7XIX (C), S BA.2 open trimer, pdb id 7XIW (D), S BA.2.75 closed trimer, pdb id 7YQU (E) and S BA.2.75 open trimer, pdb id 7YQT (F). The profiles are shown in brown, maroon and orange-colored bars for protomers A, B, and C respectively. The positions of protein stability hotspots are indicated by magenta-colored filled circles. The positions of the mutational sites for Omicron subvariants are shown in yellow-colored filled squares.

We found that the high centrality residues are assembled in tight interaction clusters with the distribution of peaks reflecting a broad interaction network in the closed states. The high centrality residues are assembled in tight interaction clusters in which the network centrality peaks often align with the hinge regions (residues 315-331, 569-572, 756-760), indicating that these key regulators of functional motions may also mediate communication in the residue interaction networks (Figure 11). The broad centrality distribution found in the closed trimers (Figure 11 A,C,E) reflects the increased number of local stabilizing clusters in the S Omicron 3RBD-down conformations in which many of the RBD mutations stabilize the inter-protomer interfaces. In the open states, we observed consolidation of local centrality peaks into three major clusters in the open form, corresponding to inter-domain borders of RBD and S2, CTD1 regions and CH residues (Figure 11B,D,F). The centrality profiles also highlighted a partial redistribution of the betweenness centrality distribution in the open forms. These observations indicated that allosteric interaction network in the open state may become more localized and operated through long-range couplings between RBD and S2 regions. Another important network signature observed in the open forms was the emergence of largest and strongest peak aligned with residues 568-572 in the CTD1 region (Figure 11).

We found that in BA.1 (Figure 11A) and BA.2.75 closed trimers (Figure 11E), the network centrality distributions are denser than in a more mobile BA.2 trimer (Figure 11C). Moreover, the significant high centrality clusters of the BA.1 and BA.2.75 trimers are aligned with the CTD1 (residues 528-591) and CTD2 (residues 592-686), while this density is weaker in the BA.2 trimer. This suggests that these regions become considerably more mobile in the closed BA.2 trimer which is consistent with the reduced stability of BA.2 and the increased heterogeneity of BA.2 states.

In general, the centrality distributions for BA.1 and BA.2.75 closed and open states are characterized by denser clusters and more pronounced peaks in the RBD, CTD and S2 regions that participate in signal transmission. The observed differences in the network centrality distributions imply that long-range allosteric interaction networks in the BA.1 and BA.2.75 are considerably stronger and broader allowing for a robust ensemble of communication pathways connecting the S1 and S2 regions. In contrast, we observed that allosteric mediating centers in a more dynamic BA.2 trimers may be more localized and concentrated in the S1-RBD core and stable S2 regions (Figure 11 C,D).

By mapping structural stability hotspots and positions of Omicron mutations, we discovered the important role of stability centers in mediating long-range interactions in the S proteins. Indeed, it could be seen that major stability centers are aligned with many local maxima of the centrality distributions, indicating a strong correspondence between stability hotspots and mediating propensities of the S protein residues (Figure 11). The network analysis has revealed a relationship between structural stability, global centrality and functional significance of hotspot residues involved in regulation. We have found that allosteric interactions in the S trimers may be mediated by modules of structurally stable residues that display high betweenness in the global interaction network. The network centrality profiles suggested that effective allosteric communications in the S trimers can be primarily provided by structurally stable residues that exhibit a significantly higher betweenness as compared to the network average. Mutational defects in functional sites may be determined by a combination of structural stability requirements and network centrality properties. The broad distribution of stable high centrality sites in BA.1 and BA.2.75 trimers suggested that allosteric communications and signal transmission in these variants may be more robust and less sensitive to mutational perturbations. Although mutations of functional residues may often result in a loss of activity, some of these changes could be tolerated and rescued by a well-connected network in which other stable network positions can offer alternative efficient routes of signal transmission. The mapping of Omicron mutational sites showed that only a fraction of the S2 sites may correspond to moderately relevant mediating centers, while Omicron RBD positions featured very small centrality (Figure 11). In network terms, a genera mechanism of allosteric interactions may involve dynamic coupling of structurally rigid and flexible residues that act cooperatively to form an ensemble of multiple communication pathways. According to the network analysis, structure stability hotspots could form the “backbone” of the communication paths that is coupled to more flexible sites, including many of the Omicron positions. Through these dynamic couplings between the stability hotspots and Omicron sites, which are determined by local and long-range interactions, mutational variants could modulate the global dynamics and allosteric ensembles of communications in the S proteins. It is possible that for more dynamic BA.2 trimers, allosteric communications could operate via a narrower “rigidity propagation path” that could be sufficiently efficient but more sensitive to mutational defects that could impair allostery. These findings support our hypothesis and provide some plausible rationale for mechanisms in which Omicron mutations can evolve to balance thermodynamic stability and conformational adaptability in order to ensure proper tradeoff between stability, binding and immune escape. The presented analysis of the S BA.1, BA.2 and BA.2.75 trimers suggested that thermodynamic stability of BA.1 and BA.2.75 variants may be intimately linked with the residue interaction network organization that allows for a broad ensemble of allosteric communications in which signaling between structural stability hotspots may be modulated by the Omicron mutational sites.

## Conclusions

In this study, we performed CG-BD molecular simulations, a comprehensive mutational scanning using two different approaches dynamic network analysis of open and closed states for the S trimers for BA.1, BA.2 and BA.2.75 Omicron variants. The results revealed differences in the dynamic and energetic patterns of the Omicron variants, allowing for a detailed characterization and analysis of the protein stability hotspots. We have shown that BA.2.75 variant can be characterized by the greater stability among the studied Omicron variants, while BA.2 trimers are characterized by a considerable mobility and are the least stable. Our findings agree with the latest experimental data and provide rationale behind the enhanced stability of the BA.2.75 trimers as one of the factors driving its increased infectivity and binding. We have found the increased number of positions in the open BA.2.275 trimer that are mutation-sensitive, indicating that BA.2.75 trimer may be more stable than BA.2 and comparable in stability to BA.1 variant. The network analysis has revealed a relationship between structural stability, global centrality and functional role of hotspot residues. The results suggested that allosteric interactions in the S trimers may be mediated by modules of structurally stable residues that display high betweenness in the global interaction network. Structure stability hotspots could form the “backbone” of the communication paths that is coupled to more flexible sites, including many of the Omicron positions. Through these dynamic couplings between the stability hotspots and Omicron sites, which are determined by local and long-range interactions, mutational variants could modulate the global dynamics and allosteric ensembles of communications in the S proteins. The results of this study highlighted the underlying mechanism of S proteins that is linked with the subtle balancing and tradeoff between protein stability, conformational adaptability and plasticity as well as immune escape pathways that collectively drive evolution of the Omicron variants and emergence of BA.2 and BA.2.75 subvariants that are markedly different in their effect on modulating conformational dynamics and protein stability.

## Materials and Methods

### Coarse-Grained Brownian Dynamics Simulations

All structures were obtained from the Protein Data Bank [112]. During structure preparation stage, protein residues in the crystal structures were inspected for missing residues and protons. Hydrogen atoms and missing residues were initially added and assigned according to the WHATIF program web interface [113]. The missing loops in the studied cryo-EM structures of the SARS-CoV-2 S protein were reconstructed and optimized using template-based loop prediction approaches ModLoop [114] and ArchPRED server [115]. The side chain rotamers were refined and optimized by SCWRL4 tool [116]. The protein structures were then optimized using atomic-level energy minimization with composite physics and knowledge-based force fields as implemented in the 3Drefine method [117]. Coarse-grained Brownian dynamics (BD) simulations have been conducted using the ProPHet (Probing Protein Heterogeneity) approach and program [81–84]. BD simulations are based on a high resolution CG protein representation [118] of the SARS-CoV-2 S Omicron trimer structures that can distinguish different residues. In this model, each amino acid is represented by one pseudo-atom at the Cα position, and two pseudo-atoms for large residues. The interactions between the pseudo-atoms are treated according to the standard elastic network model (ENM) in which the pseudo-atoms within the cut-off parameter, *R*c = 9 Å are joined by Gaussian springs with the identical spring constants of *γ* = 0.42 N m^−1^ (0.6 kcal mol^−1^ Å^−2^. The simulations use an implicit solvent representation via the diffusion and random displacement terms and hydrodynamic interactions through the diffusion tensor using the Ermak-McCammon equation of motions and hydrodynamic interactions as described in the original pioneering studies that introduced Brownian dynamics for simulations of proteins [119,120]. The stability of the SARS-CoV-2 S Omicron trimers was monitored in multiple simulations with different time steps and running times. We adopted Δ*t* = 5 fs as a time step for simulations and performed 100 independent BD simulations for each system using 100,000 BD steps at a temperature of 300 K. The CG-BD conformational ensembles were also subjected to all-atom reconstruction using PULCHRA method [121] and CG2AA tool [122] to produce atomistic models of simulation trajectories.

### Electrostatic Calculations

In the framework of continuum electrostatics, the electrostatic potential *ϕ* for biological macromolecules can be obtained by solving the Poisson–Boltzmann equation (PBE)

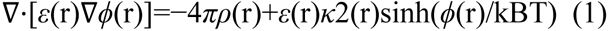

where *ϕ*(r) is the electrostatic potential, *ε*(r) is the dielectric distribution, *ρ*(r) is the charge density based on the atomic structures, *κ* is the Debye–Huckel parameter, kB is the Boltzmann constant, and T is the temperature. The electrostatic interaction potentials are computed for the averaged RBD-hACE2 conformations using the APBS-PDB2PQR software [86,87] based on the Adaptive Poisson–Boltzmann Solver (APBS) [86] and visualized using the VMD visualization tool [88]. These resources are available from the APBS/PDB2PQR website: http://www.poissonboltzmann.org/. The atomic charges and radii are assigned in this approach based on the chosen force field.

### Protein Stability Computations: Mutational Scanning and Sensitivity Analysis

We conducted mutational scanning analysis of the inter-protomer interface residues for the SARS-CoV-2 S Omicron trimers. Each residue was systematically mutated using all substitutions and corresponding protein stability changes were computed. BeAtMuSiC approach [93–95] was employed at the first stage that is based on statistical potentials describing the pairwise inter-residue distances, backbone torsion angles and solvent accessibilities, and considers the effect of the mutation on the strength of the interactions at the interface and on the overall stability of the complex. We leveraged rapid calculations based on statistical potentials to compute the ensemble-averaged binding free energy changes using equilibrium samples from simulation trajectories. The binding free energy changes were computed by averaging the results over 1,000 equilibrium samples for each of the studied systems. In the second stage, a systematic alanine scanning of the SARS-CoV-2 S Omicron inter-protomer interface residues was performed using FoldX approach [97–100]. If a free energy change between a mutant and the wild type (WT) proteins ΔΔG= ΔG (MT)-ΔG (WT) > 0, the mutation is destabilizing, while when ΔΔG <0 the respective mutation is stabilizing. We computed the average ΔΔG values using multiple samples (∼ 500) from the equilibrium ensembles using a modified FoldX protocol [99,100].

### Dynamic Network Analysis

A graph-based representation of protein structures [108,109] is used to represent residues as network nodes and the *inter-residue edges* to describe non-covalent residue interactions. The network edges that define residue connectivity are based on non-covalent interactions between residue side-chains. The residue interaction networks were constructed by incorporating the topology-based residue connectivity MD-generated maps of residues cross-correlations [110] and coevolutionary couplings between residues measured by the mutual information scores [111]. The edge lengths in the network are obtained using the generalized correlation coefficients ***R**_MI_*(***X**_i_,**X**_j_*) associated with the dynamic correlation and mutual information shared by each pair of residues. The length (i.e. weight) *w_ij_* = −log[***R**_MI_*(***X**_i_,**X**_j_*)] of the edge that connects nodes *i* and *j* is defined as the element of a matrix measuring the generalized correlation coefficient ***R**_MI_*(***X**_i_,**X**_j_*) as between residue fluctuations in structural and coevolutionary dimensions. Network edges were weighted for residue pairs with ***R**_MI_*(***X**_i_,**X**_j_*) > 0.5 in at least one independent simulation.

The Residue Interaction Network Generator (RING) program was employed for the initial generation of residue interaction networks based on the single structure [123] and the conformational ensemble [124] where edges have an associated weight reflecting the frequency in which the interaction present in the conformational ensemble. The residue interaction network files in xml format were obtained for all structures using RING v3.0 webserver freely available at https://ring.biocomputingup.it/submit. Network graph calculations were performed using the python package NetworkX [125]. Using the constructed protein structure networks, we computed the residue-based betweenness parameter. The short path betweenness of residue *i* is defined to be the sum of the fraction of shortest paths between all pairs of residues that pass through residue *i*:

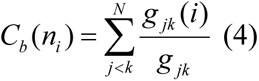

Where *g _jk_* denotes the number of shortest geodesics paths connecting *j* and *k*, and *g _jk_* (*i*) is the number of shortest paths between residues *j* and *k* passing through the node *n_i_*. Residues with high occurrence in the shortest paths connecting all residue pairs have a higher betweenness values. For each node *n*, the betweenness value is normalized by the number of node pairs excluding *n* given as (*N* - 1)(*N* - 2) / 2, where *N* is the total number of nodes in the connected component that node *n* belongs to. The normalized short path betweenness of residue *i* can be expressed as follows:

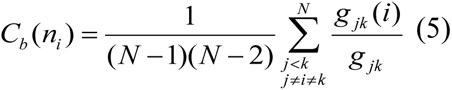

*g _jk_* is the number of shortest paths between residues *j* and k; *g _jk_* (*i*) is the fraction of these shortest paths that pass through residue *i*

## Supplementary Materials

Supplementary materials can be found at www.mdpi.com/xxx/s1.

## Author Contributions

Conceptualization, G.V.; methodology, G.V.; software, G.V.; M.A.; G.G; validation, G.V.; formal analysis, G.V.; M.A.; G.G.; investigation, G.V.; resources, G.V.; M.A.; G.G.; data curation, G.V.; writing—original draft preparation, G.V.; writing—review and editing, G.V.; M.A.; G.G.; visualization, G.V.; supervision, G.V.; project administration, G.V.; funding acquisition, G.V. All authors have read and agreed to the published version of the manuscript.

## Acknowledgments

The author acknowledges support from Schmid College of Science and Technology at Chapman University for providing computing resources at the Keck Center for Science and Engineering.

