## Supplemental Figures S1-S6 for "Coarse-Grained Molecular Simulations and Ensemble-Based Mutational Profiling of Protein Stability in the Different Functional Forms of the SARS-CoV-2 Spike Trimers : Balancing Stability and Adaptability in BA.1, BA.2 and BA.2.75 Variants"

Received: date; Accepted: date; Published: date

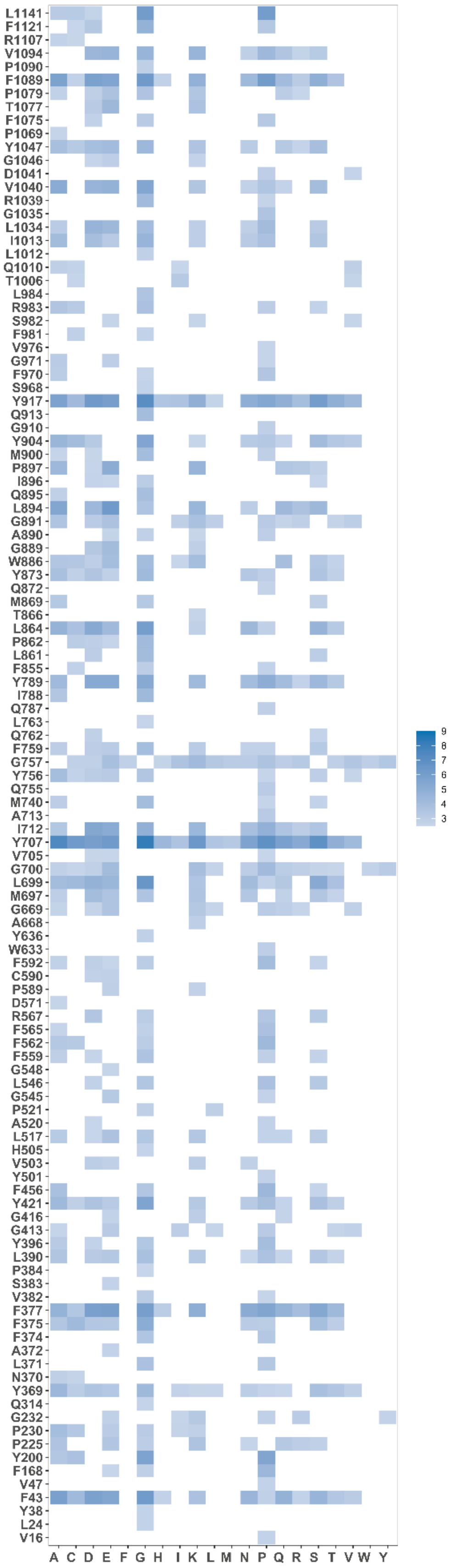


**Figure S1.** The ensemble-based dynamic mutational profiling of the S trimer inter-protomer interfacial interfaces in the S Omicron BA.1 closed trimer (pdb id 7WK2). The heatmap reports the results of systematic mutational scanning for the BA.1 closed trimer residues. For clarity of presentation, only positions in which mutations produced significant destabilization changes in the range of 2.5 kcal/mol < ΔΔG values < 9 kcal/mol are shown. To make these heatmaps more informative, the free energy changes that are outside of this range are highlighted in white color.

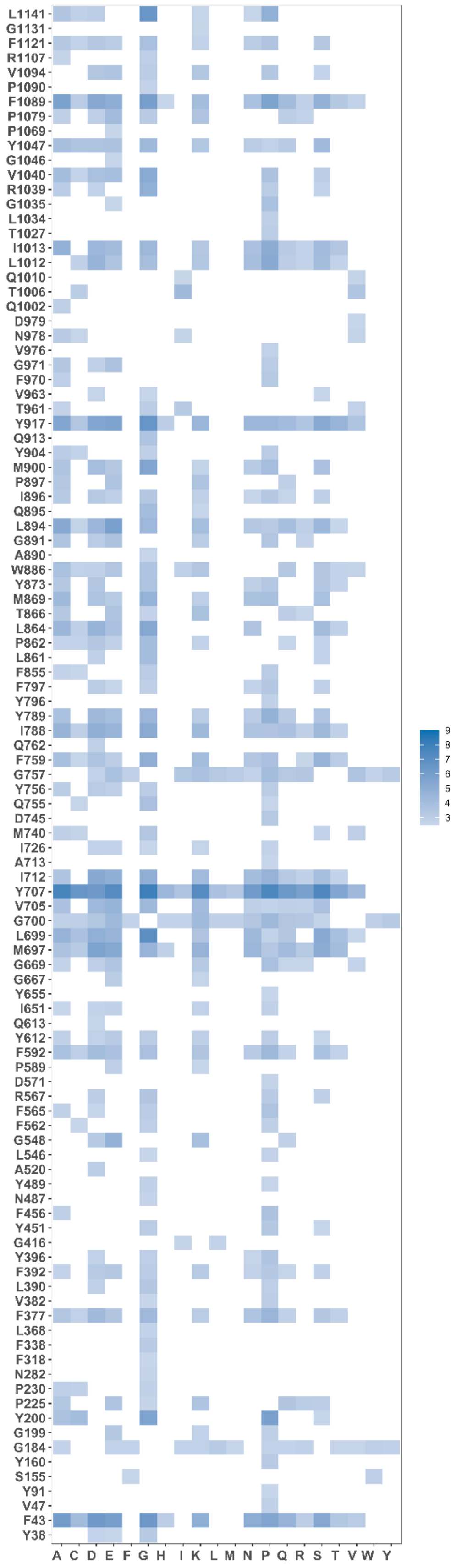

**Figure S2.** The ensemble-based dynamic mutational profiling of the S trimer inter-protomer interfacial interfaces in the S Omicron BA.1 open trimer (pdb id 7WK3). The heatmap reports the results of systematic mutational scanning for the BA.1 open trimer residues. For clarity of presentation, only positions in which mutations produced significant destabilization changes in the range of 2.5 kcal/mol < ΔΔG values < 9 kcal/mol are shown. To make these heatmaps more informative, the free energy changes that are outside of this range are highlighted in white color.

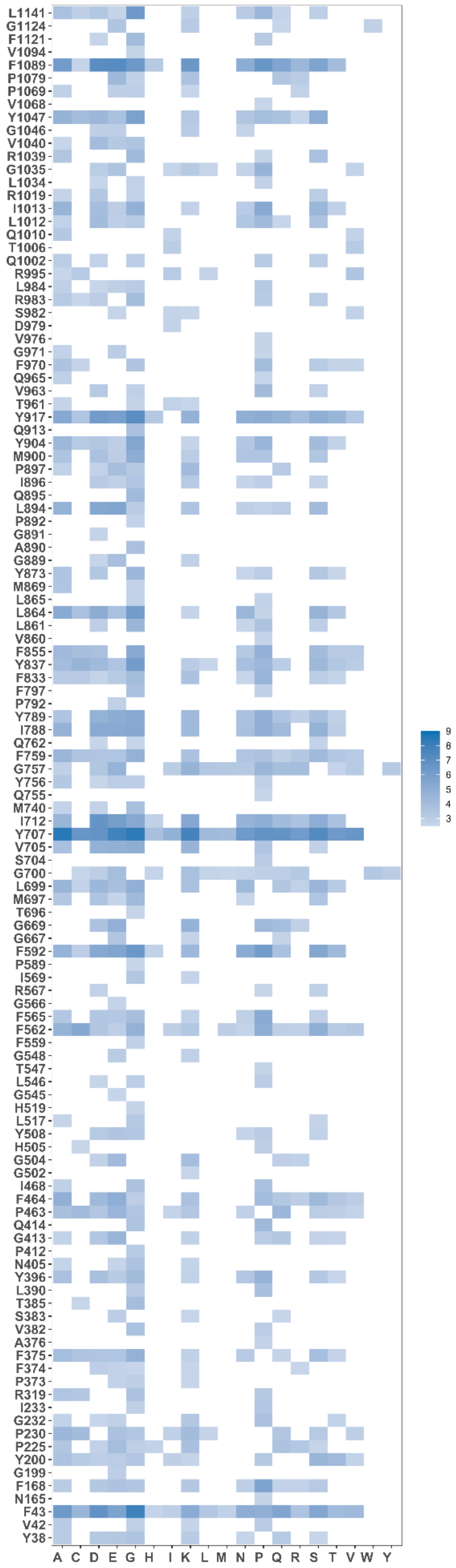

**Figure S3.** The ensemble-based dynamic mutational profiling of the S trimer inter-protomer interfacial interfaces in the S Omicron BA.2 closed trimer (pdb id 7XIX). The heatmap reports the results of systematic mutational scanning for the BA.2 closed trimer residues. For clarity of presentation, only positions in which mutations produced significant destabilization changes in the range of 2.5 kcal/mol < ΔΔG values < 9 kcal/mol are shown. To make these heatmaps more informative, the free energy changes that are outside of this range are highlighted in white color.

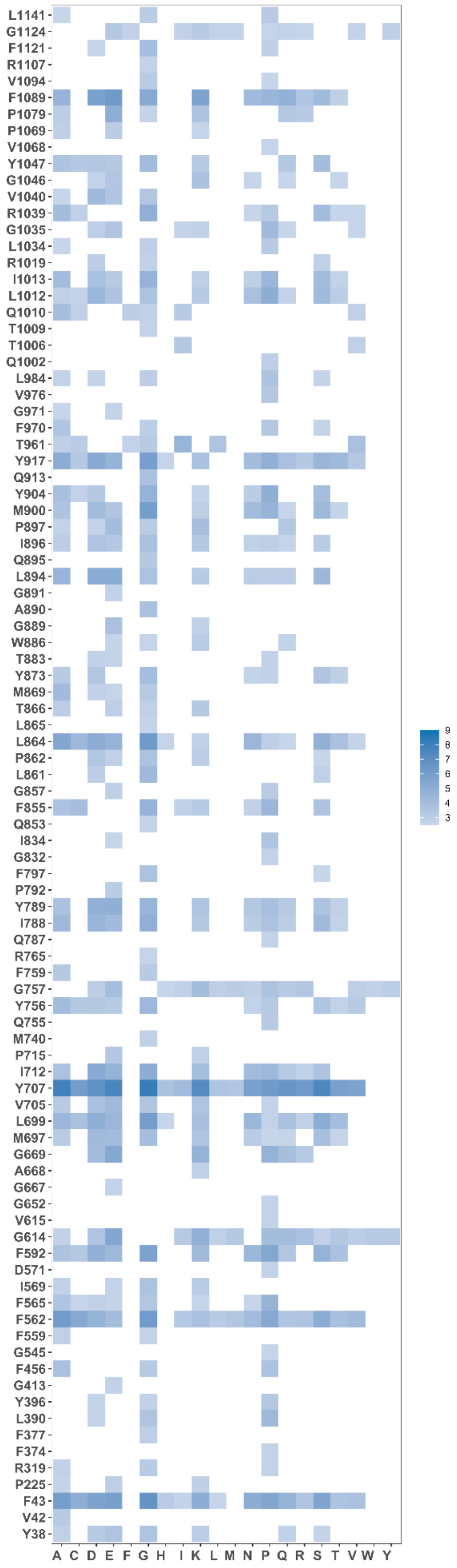


**Figure S4.** The ensemble-based dynamic mutational profiling of the S trimer inter-protomer interfacial interfaces in the S Omicron BA.2 open trimer (pdb id 7XIW). The heatmap reports the results of systematic mutational scanning for the BA.2 open trimer residues. For clarity of presentation, only positions in which mutations produced significant destabilization changes in the range of 2.5 kcal/mol < ΔΔG values < 9 kcal/mol are shown. To make these heatmaps more informative, the free energy changes that are outside of this range are highlighted in white color.

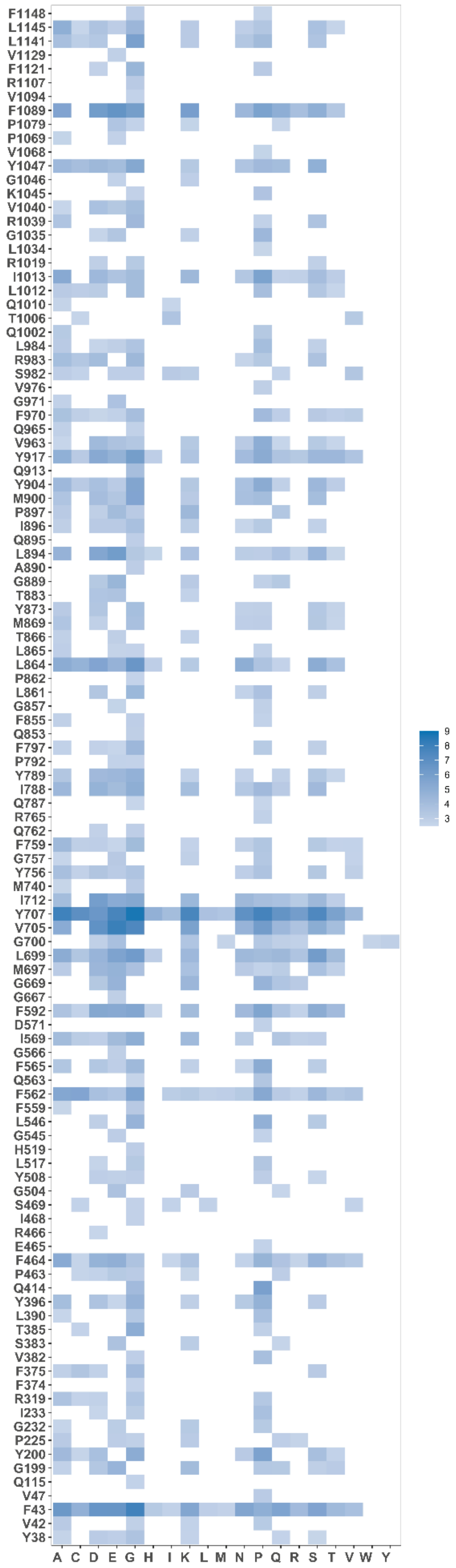


**Figure S5.** The ensemble-based dynamic mutational profiling of the S trimer inter-protomer interfacial interfaces in the S Omicron BA.2.75 closed trimer (pdb id 7YQU). The heatmap reports the results of systematic mutational scanning for the BA.2.75 closed trimer residues. For clarity of presentation, only positions in which mutations produced significant destabilization changes in the range of 2.5 kcal/mol < ΔΔG values < 9 kcal/mol are shown. To make these heatmaps more informative, the free energy changes that are outside of this range are highlighted in white color.

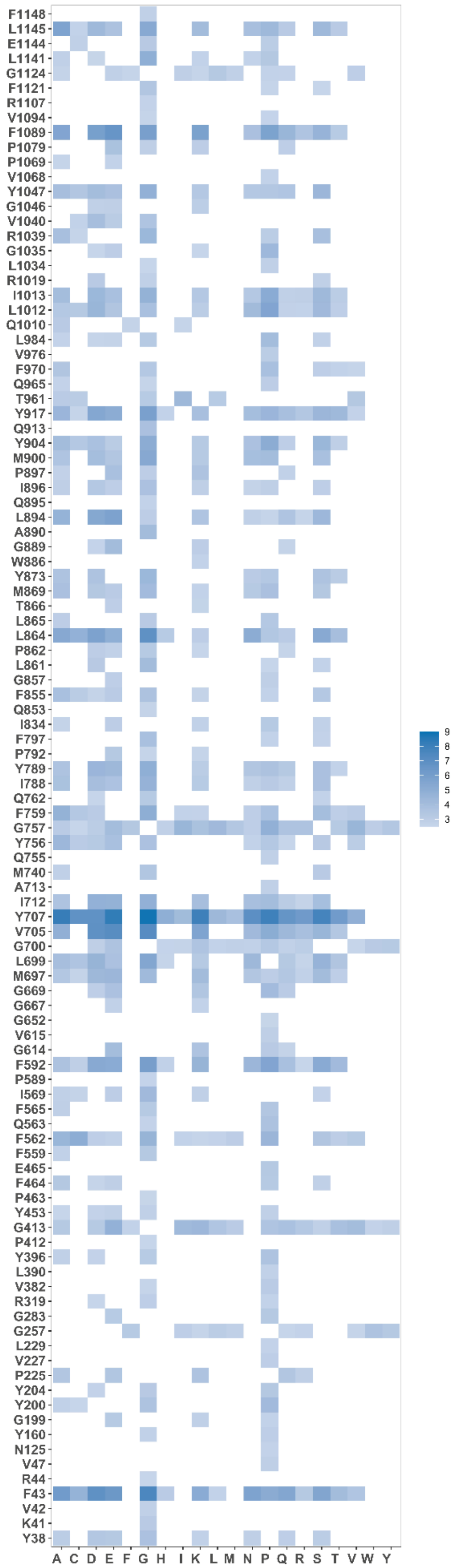


**Figure S6.** The ensemble-based dynamic mutational profiling of the S trimer inter-protomer interfacial interfaces in the S Omicron BA.2.75 open trimer (pdb id 7YQT). The heatmap reports the results of systematic mutational scanning for the BA.2.75 open trimer residues. For clarity of presentation, only positions in which mutations produced significant destabilization changes in the range of 2.5 kcal/mol < ΔΔG values < 9 kcal/mol are shown. To make these heatmaps more informative, the free energy changes that are outside of this range are highlighted in white col
